# BRIDGE-GRN: Role-Aware Bi-Tower Graph Learning with Cross-View Contrast for Directed Gene Regulatory Network Inference

**DOI:** 10.64898/2026.05.12.724562

**Authors:** Hao Chen, Wenze Ding

## Abstract

Inferring directed gene regulatory networks (GRNs) from single-cell RNA sequencing (scRNA-seq) data remains difficult because expression profiles are sparse, regulatory priors are incomplete, and experimentally supported TF–target labels are limited. To address these challenges, we propose BRIDGE-GRN, a role-aware graph learning framework that separates shared graph-context encoding from directional edge decoding. BRIDGE-GRN constructs an undirected support graph from training positive regulatory evidence, learns shared gene representations with an attention-based graph encoder, and projects them into transcription factor-role and target-role embedding spaces for asymmetric TF-to-target scoring. To improve robustness under noisy and incomplete supervision, the model aligns identity and edge-perturbed graph views through cross-view contrastive regularization. We evaluated BRIDGE-GRN across mouse benchmark settings spanning five cell types, three prior-network families, and two gene-scale settings, and further examined low-supervision transfer to target domains, architectural ablations, and biological interpretability. BRIDGE-GRN achieved consistently strong performance, outperforming or matching the strongest competing baseline in most benchmark configurations. Transfer initialization improved low-shot target-domain adaptation, while ablation analyses confirmed the importance of both role-specific bi-tower projections and contrastive regularization. Biological interpretation analyses further showed role-structured embeddings, enrichment of top-ranked predictions for external regulatory support, and coherent driver-centered regulatory modules. These results support BRIDGE-GRN as a robust, transferable, and interpretable framework for directed GRN inference from single-cell transcriptomic data.

## 1 Introduction

Reconstructing directed gene regulatory networks (GRNs) from single-cell RNA sequencing (scRNA-seq) data is a central problem for understanding cell identity, lineage progression, and cellular responses to perturbation. The rapid maturation of droplet-based profiling technologies has made it possible to measure transcriptomes for tens of thousands to hundreds of thousands of cells in a single experiment, thereby enabling increasingly fine-grained maps of tissues, developmental trajectories, and cell-state transitions [1, 2]. More recently, single-cell multi-omic measurements, functional enhancer mapping, and large pretrained models have expanded the scope of regulatory analysis, raising the possibility that regulatory structure may be inferred more accurately and transferred more effectively across datasets and biological contexts [3–9]. Despite these opportunities, many practical settings still require directed GRN inference from scRNA-seq alone, where expression matrices are sparse and noisy, cell populations are compositionally heterogeneous, prior regulatory knowledge is incomplete, and curated TF–target labels remain limited for supervision and validation [10–12].

Classical GRN inference approaches fall broadly into several families. Ensemble tree-based methods such as GENIE3 estimate directed influence through predictor importance in supervised regression models, but they were originally developed in settings closer to bulk expression data and may be sensitive to sparsity and weak signal [13]. Information-theoretic methods such as PIDC exploit multivariate dependence structures to reduce indirect associations, yet they still operate primarily on coexpression-derived statistics and do not directly encode directional asymmetry in representation space [14]. Dynamical approaches such as SCODE attempt to recover regulatory structure from pseudo-temporal variation, but they depend on trajectory assumptions that may be unstable in complex single-cell settings [15]. In parallel, motif-guided pipelines such as SCENIC integrate cis-regulatory evidence to refine target assignments, but they still rely on coexpression-based seeds and remain sensitive to the quality of prior information [16]. Benchmarking studies have therefore shown that performance varies substantially across datasets and experimental conditions, with directionality, sparsity, and class imbalance remaining persistent weaknesses across methods [11].

Recent neural and graph-based approaches have begun to address these limitations by treating GRN inference as a structured prediction problem over genes and regulatory links. Neural architectures and graph models can incorporate nonlinear feature interactions, local graph context, and prior regulatory structure more flexibly than purely correlation-based methods [17–20]. At the same time, recent progress in single-cell multi-omic integration and regulatory network reconstruction has highlighted the importance of robustness, transferability, and biological interpretability in GRN inference [21–24]. Yet an important gap remains. Many existing approaches either rely on richer multimodal inputs than standard scRNA-seq alone, or improve prediction without explicitly disentangling undirected contextual aggregation from directed edge scoring. In practice, however, directed GRN inference often requires learning from sparse scRNA-seq profiles, incomplete priors, and limited labeled edges while still producing interpretable TF-to-target predictions.

In this work, we introduce BRIDGE-GRN (*Bi-tower Encoding with Cross-View Contrast for Directed GRN Inference*), a graph representation learning framework designed specifically for this setting. Our first design principle is to separate shared context learning from directional decoding. We construct an undirected support graph from available positive TF–target pairs and learn graph-aware gene representations with a lightweight multi-head attention encoder [25]. We then project the shared representation into two role-specific spaces, one for transcription factors and one for putative targets, so that directionality is expressed through ordered TF–target pairs at decoding time. The resulting score is computed by a dot product between TF-role and target-role embeddings, which remains asymmetric because the two towers have distinct parameters. Our second design principle is to improve robustness under sparse and noisy supervision. To this end, we use a dual-view training scheme in which the encoder is exposed to an identity view and a perturbed edge-drop view of the support graph, while a node-level contrastive objective aligns the resulting representations across views. Contrastive learning has proved effective for learning invariant structure from graph augmentations [26], and here it serves as a regularizer that reduces over-reliance on any single graph realization. This is especially relevant in directed GRN inference, where false negatives, imperfect priors, and severe class imbalance can otherwise destabilize training.

These design choices yield a compact framework that learns graph-aware gene embeddings, expresses directionality through role-specific projections, and remains robust under sparse supervision. Our experiments follow established benchmarking protocols to ensure comparability and reproducibility [11]. BRIDGE-GRN makes three main contributions. First, it introduces a role-aware bi-tower framework that decouples undirected graph-context aggregation from directed TF–target link prediction. Second, it incorporates cross-view contrastive regularization to improve representation stability under support-graph perturbation and incomplete regulatory supervision. Third, it provides a comprehensive evaluation across mouse benchmark settings, low-supervision transfer analyses, architectural ablations, and biological interpretation experiments, thereby assessing not only predictive performance but also transferability, component-level contribution, and regulatory interpretability.

## 2 Methods

### 2.1 Problem Formulation

Let *X* ∈ℝ^*n* ×*d*^ denote the gene-by-cell expression matrix for *n* genes and *d* cells after preprocessing. In our implementation, each gene is standardized across cells using a z-score transformation:

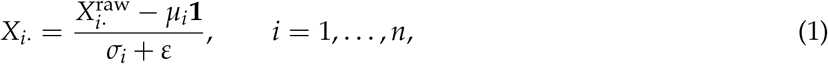

where *μ*_*i*_ and *σ*_*i*_ are the mean and standard deviation of gene *i* across cells, and *ε>* 0 is a small constant for numerical stability.

We aim to infer a directed gene regulatory network over transcription factors (TFs) and target genes. Let *τ* ⊆{1, …, *n*} denote the index set of TFs and let 𝒢⊆{1, …, *n*} denote the candidate target set. Supervision is provided as labeled TF–target pairs

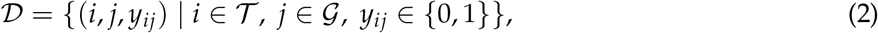

where *y*_*ij*_ = 1 indicates a known regulatory edge from TF *i* to target *j*, and *y*_*ij*_ = 0 indicates a sampled negative pair.

For each ordered pair (*i, j*), the model outputs a logit score *s*(*i* ⟶*j*) ∈ℝand a probability

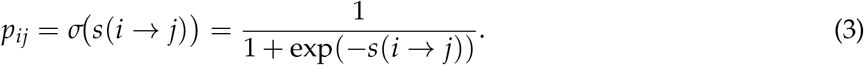

#### Support graph for message passing

We construct a support graph from positive training labels only. For each positive TF–target pair in the training set, we add an undirected edge to the support graph:

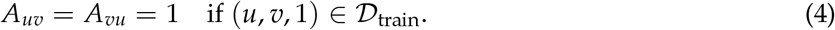

Negative pairs do not contribute to the support graph. The support graph is constructed from training positives only, thereby avoiding information leakage from validation or test edges. In implementation, nodes without retained neighbors are handled by preserving their own transformed features during message passing, preventing empty-neighborhood attention from inducing spurious aggregation.

### 2.2 Dual-View Graph Augmentation

At each forward pass, the model generates two graph views that share the same node features *X* but differ in their edge sets. The first view is the identity view, which keeps the support graph unchanged. The second view is obtained by independently removing edges from the support graph representation with probability *p*_*e*_.

Let *E* denote the set of edges stored in the graph index representation. For each edge *e* ∈*E*, an independent Bernoulli variable *m*_*e*_ ~Bernoulli(1 −*p*_*e*_) is sampled, and the edge is retained if *m*_*e*_ = 1. Thus the second view is formed by

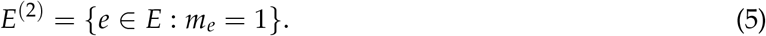

In the implementation, if all edges are removed by random sampling, one edge is randomly restored so that the perturbed view is never empty.

Because edge removal is applied independently to entries in the graph index representation, the perturbed view is not forced to remain exactly symmetric even when the original support graph is undirected.

### 2.3 Attention-Based Graph Encoder

Let *H*^(0)^ = *X*. We use a two-layer multi-head graph attention encoder. In each attention head, node features are first linearly transformed:

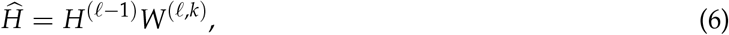

For each head, attention logits are computed for all ordered node pairs:

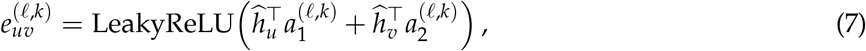

where 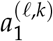 and 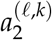 are learnable attention vectors. In the implementation, these logits are computed as a dense matrix over all node pairs and then masked by the adjacency matrix of the current view. Specifically,

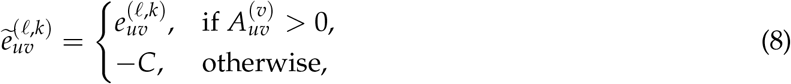

where *C* is a large positive constant. Attention coefficients are then obtained by row-wise softmax:

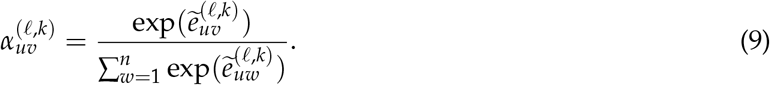

The output of each head is

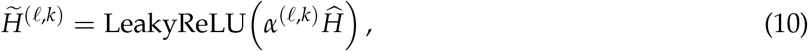

followed by row-wise *ω*_2_ normalization. In the current implementation, dropout is not applied within the attention layer itself.

For the first encoder layer, multiple heads are fused either by concatenation or averaging. In the main setting used in our training scripts, the first-layer outputs are concatenated:

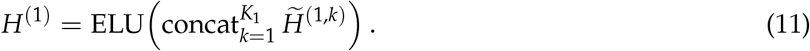

For the second encoder layer, the outputs of all heads are averaged:

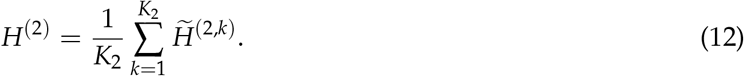

The matrix *H*^(2)^ is the pre-tower node representation used both for directed edge scoring and for contrastive regularization.

### 2.4 Role-Specific Bi-Tower Projections

To encode directionality, we map the shared representation *H*^(2)^ into two role-specific embedding spaces, one for TFs and one for targets. Each tower is implemented as a two-layer multilayer perceptron with LeakyReLU activations and dropout between the two linear layers.

For the TF tower,

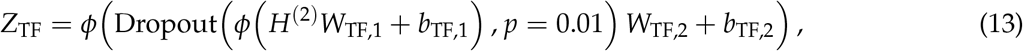

and for the target tower,

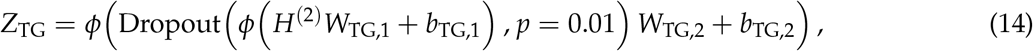

where *ϕ* (·) denotes LeakyReLU. The resulting matrices satisfy

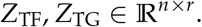

### 2.5 Directional Decoder

For an ordered TF–target pair (*i, j*), we select the TF-role embedding of node *i* and the target-role embedding of node *j*, and compute a scalar score. In the main model, we use a directional dot-product decoder:

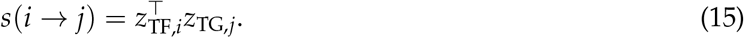

Directionality arises because the two towers have different parameters, so in general

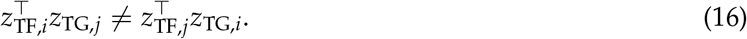

Our implementation also includes cosine-similarity and bilinear decoders as optional alternatives for comparison or ablation, but the default configuration in the main training script uses the directional dot-product decoder.

### 2.6 Supervised Link Reconstruction Loss

Let ℬ ⊂ 𝒟_train_ be a minibatch of labeled TF–target pairs. For the two graph views, the encoder produces logits *s*^(1)^(*i* →*j*) and *s*^(2)^(*i* →*j*). The supervised loss is the sum of binary cross-entropies on the sigmoid-transformed logits from both views:

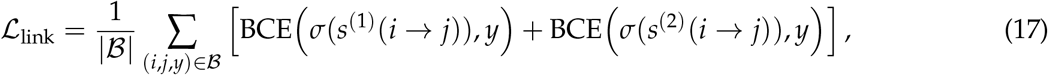

where

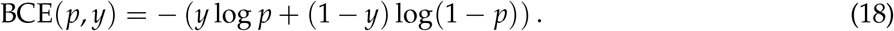

### 2.7 Cross-View Contrastive Regularization

To improve robustness to edge perturbation, we apply a symmetric InfoNCE loss between the pre-tower representations from the two views. In the implementation, this regularization is applied only to the set of TF nodes, using the complete TF index list provided by the dataset rather than only the TFs appearing in the current minibatch.

Let 𝒯denote the full TF index set, and let

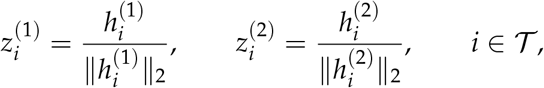

where 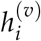 is the *i*th row of *H*^(2)^ under view *v*. We define the cosine-based similarity matrix

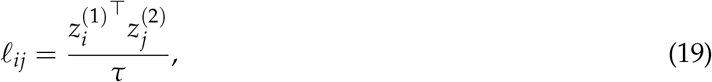

where *τ>* 0 is the temperature parameter. For the first direction, the InfoNCE loss is

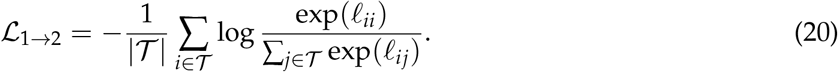

The reverse-direction loss ℒ_2→1_ is defined analogously by swapping the two views. The final contrastive loss is their average:

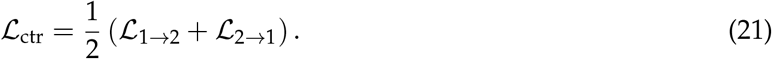

In our implementation, the temperature is fixed at *τ* = 0.2.

### 2.8 Overall Objective and Optimization

The full training objective combines supervised link reconstruction and cross-view contrastive regularization:

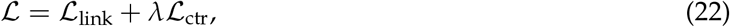

where *λ>* 0 controls the contribution of the contrastive term. In the default training configuration, we use *λ* = 0.5. All parameters are optimized end-to-end using Adam. In the implementation used for the reported experiments, the learning rate is initialized at 3 ×10^−3^ and updated with a StepLR scheduler with decay factor 0.99 after each epoch. Training is performed using the same fixed training schedule across compared model variants, and model selection is based on validation AUROC.

## 3 Results

### 3.1 Overview of BRIDGE-GRN and study design

Directed gene regulatory network (GRN) inference from single-cell RNA sequencing data requires the recovery of directed TF–target relationships from sparse expression profiles, incomplete regulatory priors, and limited experimentally supported labels. As summarized in Figure 1, BRIDGE-GRN was designed to address these challenges by separating shared graph-context encoding from role-specific directional decoding. The model takes scRNA-seq-derived gene expression profiles and training regulatory evidence as inputs, constructs an undirected support graph from positive TF–target evidence, and learns shared graph-context representations through a graph encoder. These shared representations are then projected into transcription factor-role and target-role embedding spaces, allowing asymmetric TF-to-target scoring for directed regulatory edge prediction. To reduce over-dependence on a single imperfect graph realization, BRIDGE-GRN further aligns representations from identity and edge-perturbed graph views through cross-view contrastive regularization.

**Figure 1:**
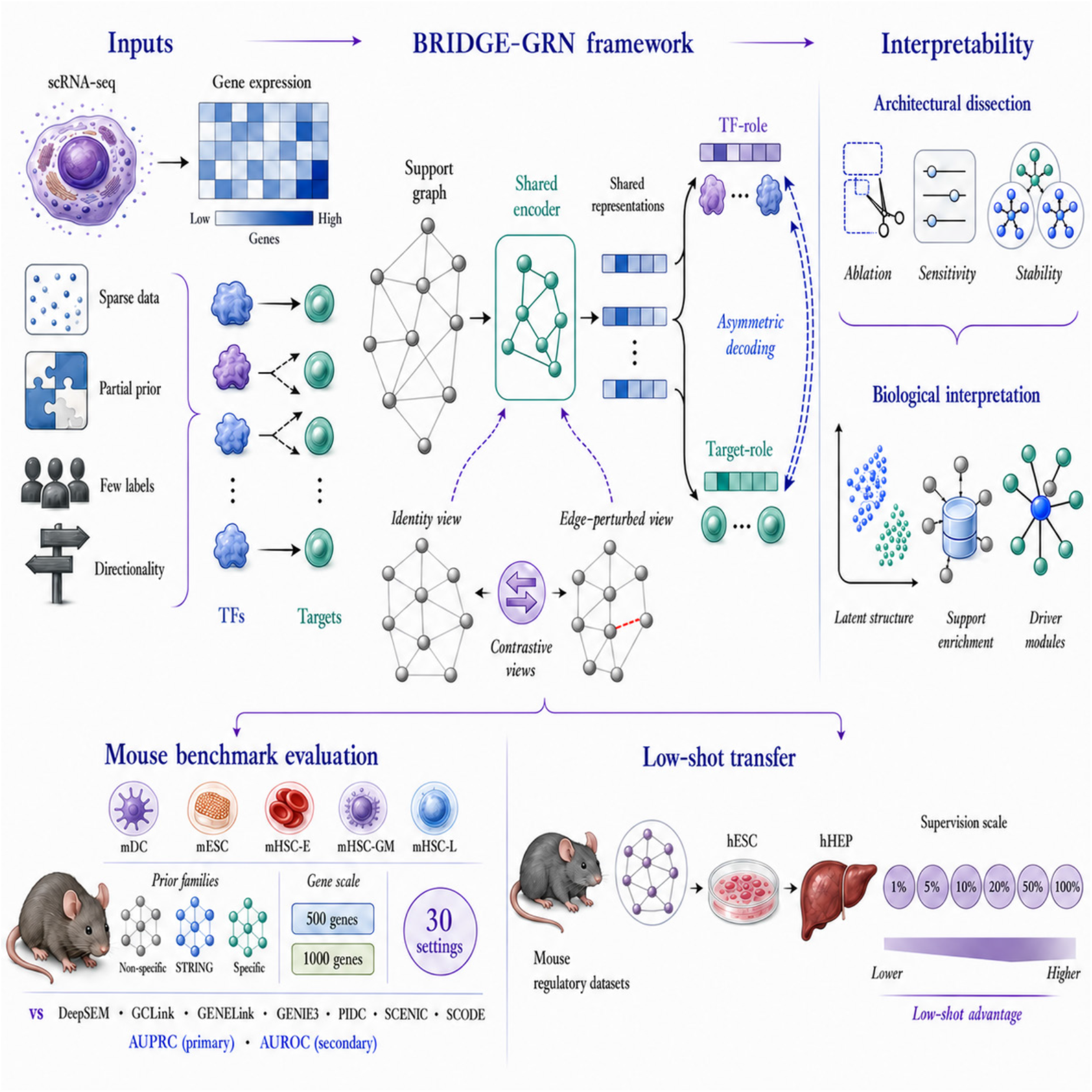
Overview of BRIDGE-GRN and study design. The figure summarizes the input data, model framework, benchmark evaluation, transfer-learning design, and interpretability analyses used to evaluate BRIDGE-GRN. Inputs include scRNA-seq gene expression profiles and regulatory evidence under common challenges of sparse expression measurements, incomplete prior knowledge, limited regulatory labels, and directional asymmetry. BRIDGE-GRN constructs an undirected support graph, learns shared graph-context representations, projects them into TF-role and target-role embedding spaces, and performs asymmetric TF-to-target decoding. Identity and edge-perturbed graph views are aligned through cross-view contrastive regularization. The empirical evaluation includes mouse benchmark analyses across cell types, prior-network families, and gene-scale settings;low-supervision transfer from source regulatory datasets to target domains including hESC and hHEP;and interpretability analyses covering architectural dissection, perturbation sensitivity, stability analysis, role-specific latent organization, external support enrichment, and driver-centered regulatory modules.

The empirical evaluation was organized into four modules, as summarized in Supplementary Table S1. First, we evaluated predictive performance across mouse benchmark settings spanning five cell types, three prior-network families, and two gene-scale settings. Second, we examined transferability using source-domain pretraining followed by low-supervision adaptation to target domains, including human target-domain settings such as hESC and hHEP. Third, we performed architectural dissection to identify the contribution of core model components, including the role-specific bi-tower structure, contrastive regularization, perturbation sensitivity, and repeated-run stability. Fourth, we examined biological interpretability through role-specific latent organization, external support enrichment, and driver-centered regulatory modules. The shared preprocessing, support-graph construction, split strategy, transfer-learning settings, and evaluation metrics are summarized in Supplementary Table S2.

### 3.2 BRIDGE-GRN generalizes across priors, biological contexts, and species

We first asked whether BRIDGE-GRN provides a stable advantage across heterogeneous benchmark settings, rather than performing well only in a small number of favorable cases. The overall evaluation framework, dataset organization, preprocessing conventions, split strategy, and metric definitions are summarized in Supplementary Tables S1–S2. To keep the within-benchmark evaluation distinct from transfer adaptation, we first focused on mouse benchmark settings, including mDC, mESC, mHSC-E, mHSC-GM, and mHSC-L, across three prior-network families and two gene-scale settings. This yielded 30 mouse benchmark configurations. BRIDGE-GRN was compared with representative baselines covering major GRN inference strategies, including DeepSEM, GCLink, GENELink, GENIE3, PIDC, SCENIC, and SCODE. AUPRC was used as the primary metric because regulatory positives are rare, and AUROC was used as a complementary measure of global ranking quality.

Across the mouse benchmark collection, BRIDGE-GRN achieved higher AUPRC than the strongest competing baseline in 29 of 30 settings, with one slight deficit in the STRING prior setting for mHSC-GM at the 1000-gene scale (Figure 2a–c; Supplementary Figures S1–S2 and Supplementary Tables S3–S5). The median absolute AUPRC gain across the 30 mouse settings was +0.0745. When stratified by prior-network family, the median AUPRC gains were +0.036 for Non-Specific priors, +0.095 for STRING priors, and +0.0815 for Specific priors. This result indicates that the model’s advantage was not restricted to curated priors, but remained visible under weaker and noisier prior-network conditions.

**Figure 2:**
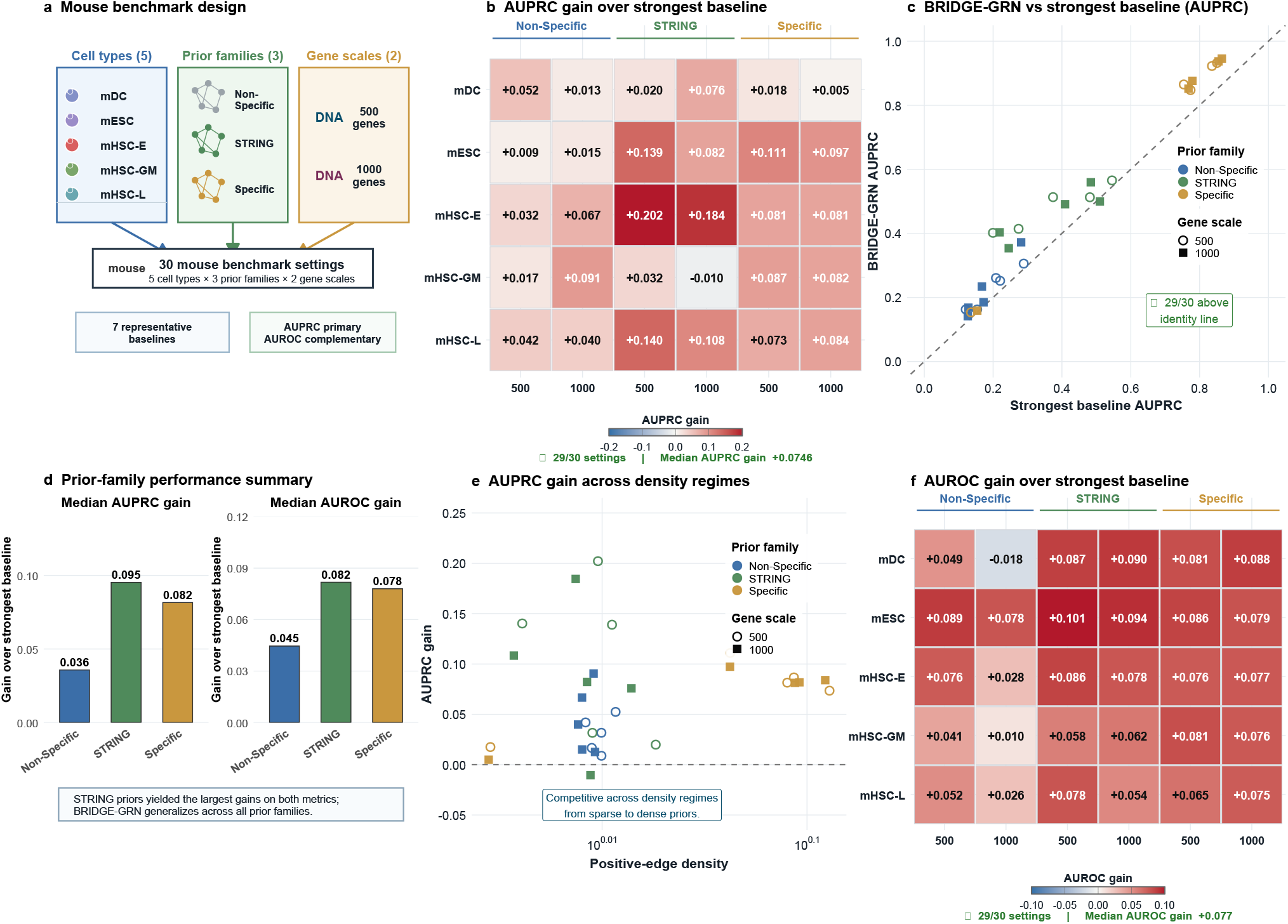
BRIDGE-GRN shows robust generalization across mouse benchmark settings with varying prior quality, biological context, and gene scale. **a**, Overview of the mouse benchmark design, including five mouse cell types, three prior-network families, and two gene-scale settings, yielding 30 benchmark configurations. **b**, Heatmap of AUPRC gains of BRIDGE-GRN over the strongest competing baseline across all mouse benchmark configurations. Values indicate absolute AUPRC differences between BRIDGE-GRN and the best-performing baseline in each setting. **c**, Paired AUPRC comparison between BRIDGE-GRN and the strongest baseline for each mouse benchmark setting. The dashed line denotes equal performance. **d**, Prior-family summary of median AUPRC and AUROC gains, showing performance improvements under Non-Specific, STRING, and Specific prior networks. **e**, Relationship between AUPRC gain and positive-edge density, illustrating that BRIDGE-GRN remains competitive across sparse and denser prior-network regimes. **f**, Heatmap of AUROC gains of BRIDGE-GRN over the strongest competing baseline across the same mouse benchmark configurations. Values indicate absolute AUROC differences.

A similar finding was observed for AUROC. BRIDGE-GRN improved over the strongest competing baseline in 29 of 30 mouse benchmark settings, with a median AUROC gain of +0.077 (Figure 2d,f; Supplementary Figure S3 and Supplementary Tables S4–S6). The corresponding prior-stratified median AUROC gains were +0.045 under Non-Specific priors, +0.082 under STRING priors, and +0.078 under Specific priors. The paired comparison in Figure 2c shows that most benchmark points lie above the identity line, confirming that the performance gain was broadly distributed rather than being driven by a small number of high-performing datasets. In addition, the gain-versus-density analysis showed that BRIDGE-GRN remained competitive across the range of positive-edge densities represented in the benchmark (Figure 2e;Supplementary Table S7), suggesting that the model was not narrowly tuned to a single graph-density regime.

We next evaluated whether BRIDGE-GRN could generalize beyond the original mouse benchmark distribution. In the transfer analysis, the model was first pretrained on source-domain regulatory datasets and then fine-tuned on target domains under limited supervision (Figure 3a). In the human target-domain subset, transfer initialization improved mean AUPRC from 0.301 to 0.465 at 1%target supervision, from 0.337 to 0.501 at 5%, and from 0.428 to 0.542 at 10%. The corresponding AUROC values increased from 0.452 to 0.750, from 0.411 to 0.783, and from 0.612 to 0.800, respectively (Figure 3b–c; Supplementary Figures S4–S7 and Supplementary Table S9). These improvements were strongest in the few-shot regime and narrowed as target-domain supervision increased, consistent with the expectation that pretrained regulatory representations are most useful when labeled target-domain edges are scarce.

**Figure 3:**
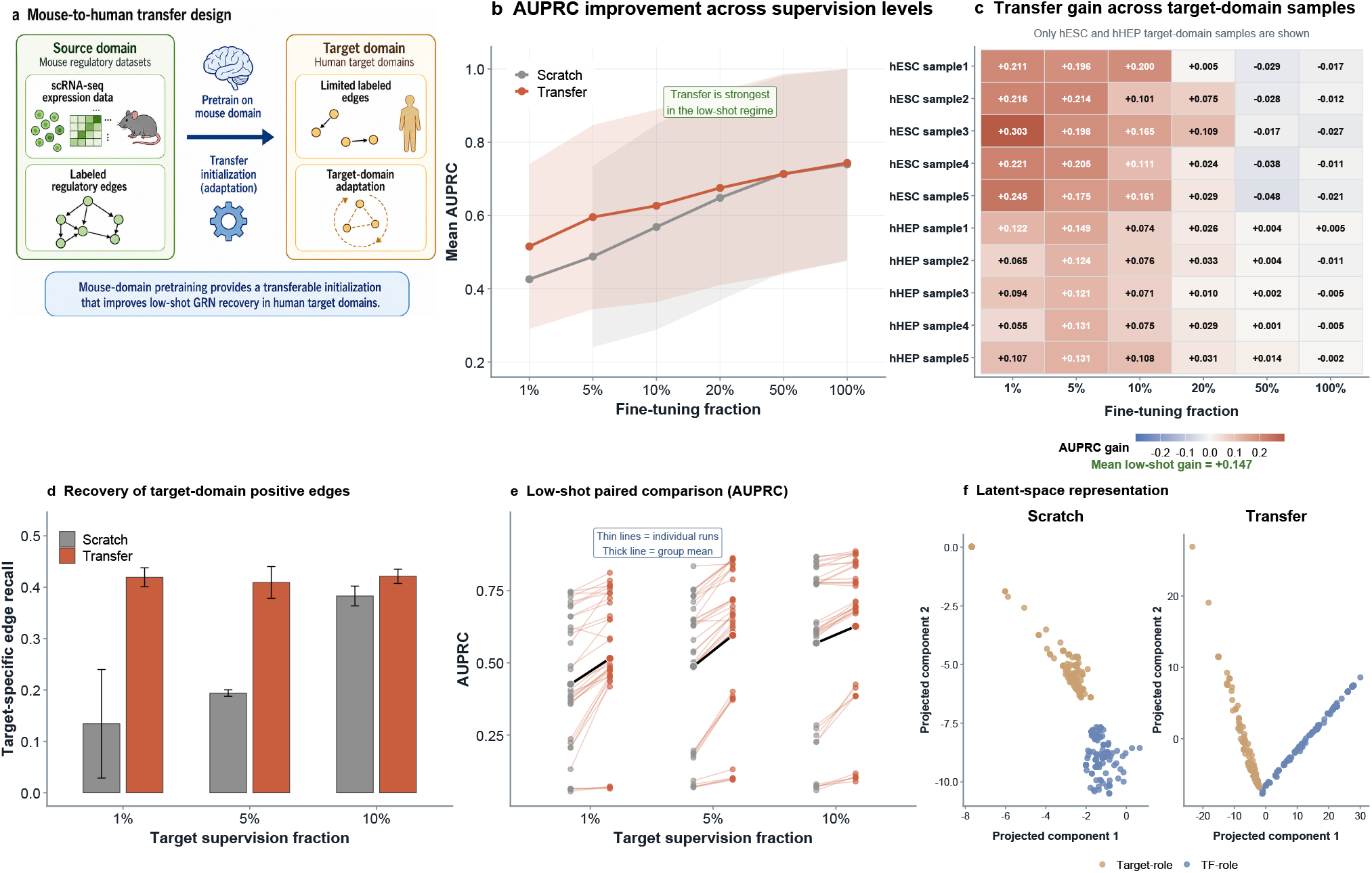
BRIDGE-GRN transfers to human target domains under limited supervision. **a**, Mouse-to-human transfer design. The model is pretrained on mouse regulatory datasets and adapted to human target domains with limited labeled edges. **b**, AUPRC curves comparing scratch training and transfer initialization across target-domain fine-tuning fractions. **c**, Transfer gain heatmap across human target-domain samples and supervision fractions. **d**, Recovery of target-domain positive regulatory edges under low supervision. **e**, Low-shot paired performance comparison between scratch and transfer initialization. **f**, Representative latent-space reorganization under scratch training and transfer initialization.

The broader transfer analysis showed the same low-supervision finding across target domains. Across all evaluated target-domain runs, mean AUPRC increased from 0.426 to 0.515 at 1%supervision, from 0.488 to 0.596 at 5%, and from 0.569 to 0.627 at 10%. AUROC increased from 0.548 to 0.719, from 0.561 to 0.793, and from 0.687 to 0.806 at the same supervision fractions (Figure 3b–e; Supplementary Figures S4–S7 and Supplementary Tables S8–S10). Transfer initialization also improved recovery of target-domain positive regulatory edges: target-specific edge recall increased from 0.134 to 0.419 at 1%supervision, from 0.194 to 0.409 at 5%, and from 0.383 to 0.421 at 10%(Figure 3d; Supplementary Figure S8 and Supplementary Table S11). Representative latent-space analyses further showed that transfer initialization reorganized role-specific embeddings relative to scratch training (Figure 3f; Supplementary Figures S9–S10 and Supplementary Table S12). Thus, transfer did not merely improve global ranking metrics, but also enhanced recovery of target-domain regulatory structure under sparse supervision.

The benchmark and transfer results support two levels of generalization. Within the mouse benchmark framework, BRIDGE-GRN remained robust across prior-network qualities, biological contexts, and gene-scale settings. Under the more difficult transfer setting, pretrained BRIDGE-GRN representations improved low-shot adaptation to target domains, including human regulatory contexts. These findings suggest that the model learns regulatory representations that are useful beyond a single dataset or prior condition.

### 3.3 Architectural dissection identifies the key drivers of performance

We next dissected the architecture to determine which components were responsible for the observed performance gains. BRIDGE-GRN combines graph-context encoding, role-specific directional factorization, cross-view regularization, and a simple directional decoder. Its improvement could therefore arise from several sources. We compared the full model with targeted ablations that removed the two central design elements: the role-specific bi-tower structure and the cross-view contrastive objective. We also examined secondary architectural choices, including decoder formulation, embedding dimension, fusion strategy, and perturbation strength.

The core ablation results showed that both central design elements were necessary for optimal performance (Figure 4a;Supplementary Tables S13–S14). Removing the cross-view contrastive objective reduced mean AUPRC from 0.876 to 0.857, while replacing the role-specific bi-tower with a single shared tower reduced mean AUPRC from 0.876 to 0.844. The latter produced the larger performance drop, indicating that explicit separation of TF-role and target-role embeddings is a major contributor to directed edge recovery. Paired comparisons across the matched repeated runs further supported this finding, with the full model outperforming the variant without contrastive regularization and the single-tower variant. These results show that the gain of BRIDGE-GRN is not attributable to the attention backbone alone, but depends on the role-aware directional factorization introduced after shared graph-context encoding.

**Figure 4:**
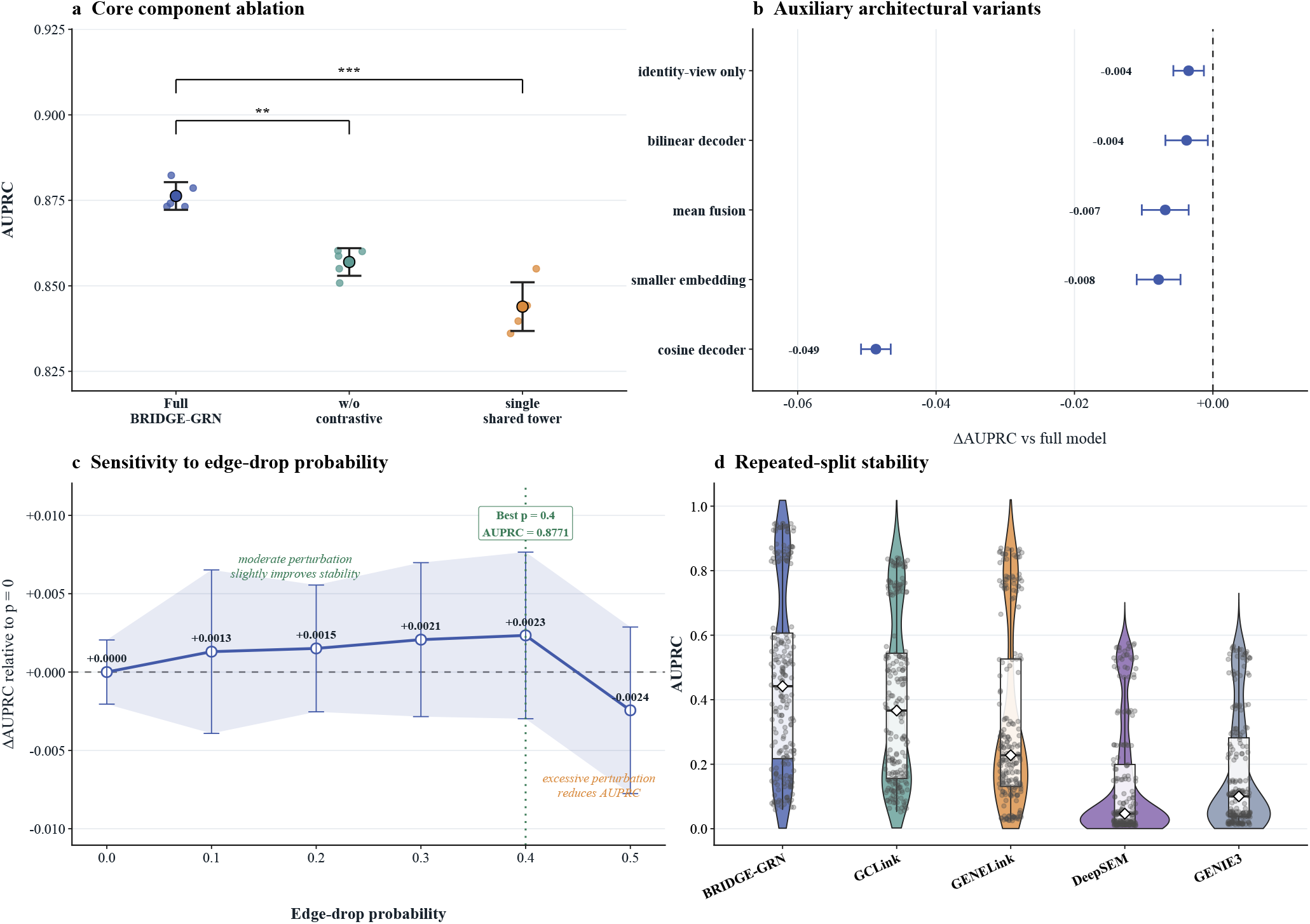
Architectural dissection identifies the key drivers of BRIDGE-GRN performance. **a**, Core component ablation comparing the full model with variants without contrastive regularization and without role-specific bi-tower projections. **b**, Effect-size summary of auxiliary architectural variants, including decoder formulation, fusion strategy, embedding dimension, and identity-view-only training. Values indicate ΔAUPRC relative to the full model. **c**, Sensitivity to the edge-drop probability used for graph-view perturbation, shown as ΔAUPRC relative to the no-perturbation setting. **d**, Stability across repeated benchmark runs compared with representative graph-based and classical baselines.

Secondary design choices modulated performance but did not explain the main gain (Figure 4b;Supplementary Table S13). The directional dot-product decoder performed comparably to the bilinear decoder while avoiding decoder-specific parameterization, with the bilinear variant showing only a small AUPRC decrease of 0.004 relative to the full model. In contrast, cosine similarity produced a much larger AUPRC decrease of 0.049, suggesting that a purely normalized similarity measure is insufficient for directed GRN scoring. Other modifications also produced smaller reductions: identity-view-only training decreased AUPRC by 0.004, mean fusion instead of concatenation decreased AUPRC by 0.007, and reducing the embedding dimension decreased AUPRC by 0.008. These changes affected performance but did not alter the overall conclusion that the model’s main advantage is driven by role-specific directional decomposition and cross-view regularization.

The perturbation-sensitivity analysis further supported the contrastive design (Figure 4c;Supplementary Table S14). Because the absolute AUPRC differences across edge-drop probabilities were small, we visualized the change relative to the no-perturbation setting (*p* = 0). Moderate edge-drop probabilities slightly improved or maintained performance, with the best setting at *p* = 0.4 increasing AUPRC from 0.8748 to 0.8771. In contrast, stronger perturbation at *p* = 0.5 reduced AUPRC to 0.8723. This result is consistent with the intended role of the perturbed view: mild structural perturbation regularizes the encoder and discourages over-reliance on the observed support graph, whereas overly strong perturbation removes useful relational information.

Across repeated benchmark runs, BRIDGE-GRN also maintained a higher AUPRC distribution than the representative graph-based and classical baselines considered in the stability analysis (Figure 4d). The violin distributions show that BRIDGE-GRN achieved consistently strong performance across heterogeneous benchmark settings, while several competing methods showed lower central performance or weaker high-performing regimes. These analyses identify the role-specific bi-tower and cross-view contrastive regularization as the principal drivers of BRIDGE-GRN performance, with auxiliary design choices refining but not replacing these core mechanisms.

### 3.4 BRIDGE-GRN recovers biologically coherent and interpretable regulatory structure

Beyond predictive accuracy, a useful GRN inference model should recover regulatory structure that can be interpreted biologically. We therefore examined BRIDGE-GRN outputs at three levels: role-specific latent representations, edge-level external support, and driver-centered regulatory modules. This analysis was intended to determine whether the model learns organized regulatory programs rather than only improving benchmark ranking scores.

First, the learned TF-role and target-role embeddings separated in latent space (Figure 5a; Supplementary Figure S11). This separation is consistent with the architectural design of BRIDGE-GRN, in which directionality is not imposed during undirected context aggregation but is instead introduced through role-specific projections. Quantitatively, the two role spaces showed distinct coordinate distributions and clustering structure (Supplementary Figure S12 and Supplementary Table S15). The role-specific embeddings were therefore related but not interchangeable, supporting the interpretation that the bi-tower architecture learns different regulatory roles for the same set of genes.

**Figure 5:**
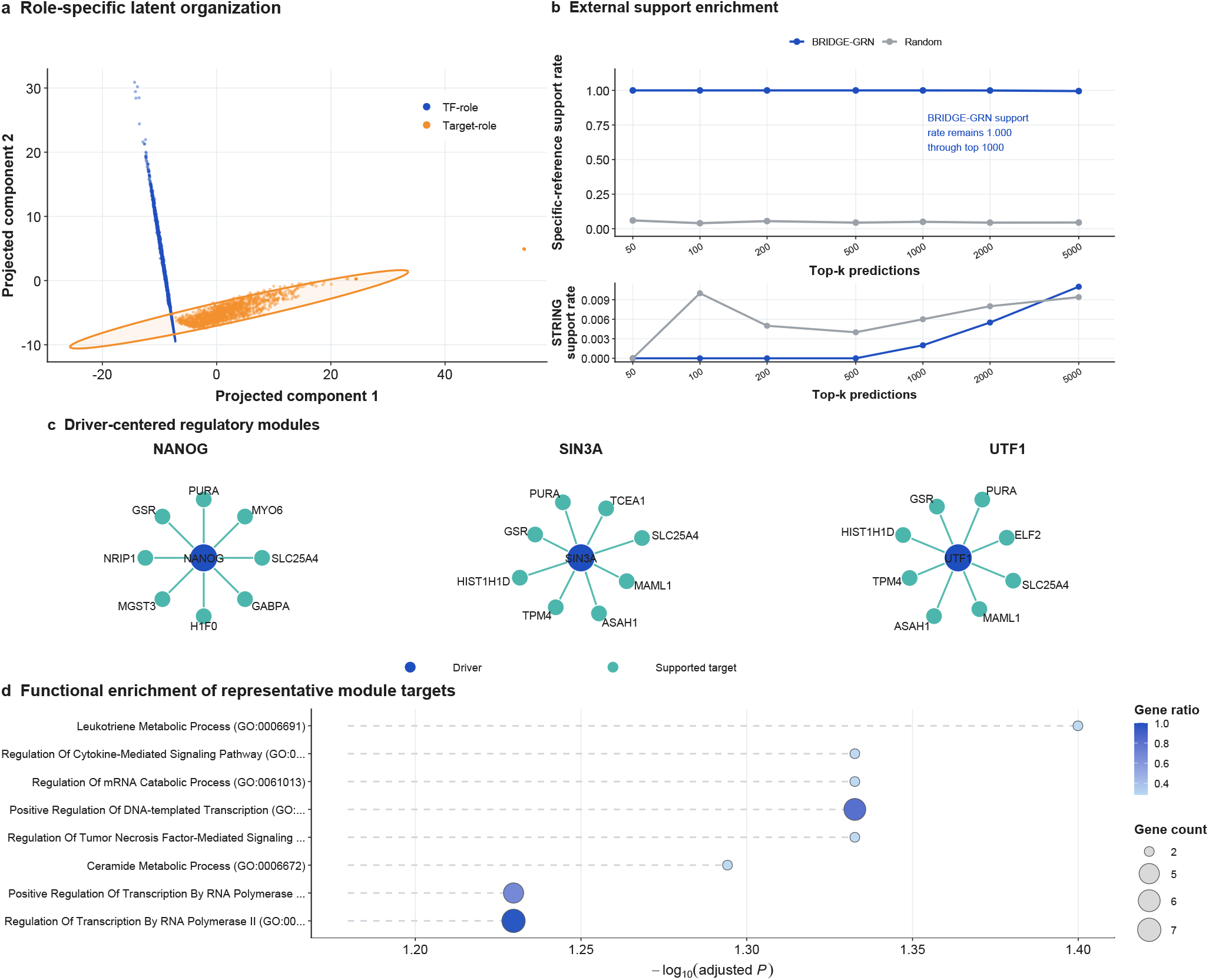
BRIDGE-GRN recovers biologically coherent and interpretable regulatory structure. **a**, Projection of TF-role and target-role embeddings showing role-specific latent organization. **b**, External support rate of top-ranked predictions compared with random ranking under specific-reference and STRING-overlap definitions. **c**, Driver-centered regulatory modules inferred by BRIDGE-GRN, including NANOG-, SIN3A-, and UTF1-centered modules. **d**, Functional enrichment of representative module targets.

Second, top-ranked predicted edges were enriched for external regulatory support (Figure 5b; Supplementary Figure S13 and Supplementary Tables S16–S17). Under the specific-reference support definition, the support rate for BRIDGE-GRN remained at 1.000 through the top 1000 predictions, whereas the corresponding random baseline remained near 0.04–0.06. Support under STRING overlap was much weaker, with the mean BRIDGE-GRN support rate remaining close to zero across the evaluated top-*k* thresholds. This result indicates that the strongest external support was obtained under the more curated reference definition, while broader interaction overlap was comparatively sparse. The asymmetry suggests that BRIDGE-GRN preferentially ranks edges supported by specific regulatory evidence, while also identifying high-confidence regulatory candidates that may not be fully represented in broader interaction resources.

Third, high-confidence predictions organized into compact driver-centered modules rather than diffuse sets of isolated edges. In the representative mESC analysis, BRIDGE-GRN recovered regulatory modules centered on NANOG, SIN3A, and UTF1 (Figure 5c;Supplementary Table S18). The NANOG-centered module included supported outgoing regulatory links to targets such as PURA, GSR, NRIP1, MGST3, H1F0, GABPA, SLC25A4, and MYO6. Similarly, the SIN3A- and UTF1-centered modules showed compact sets of supported target links. These driver-centered structures indicate that the model output can be summarized as interpretable regulatory programs rather than only as an unstructured ranked edge list.

Finally, the targets within representative driver-centered modules showed functional enrichment for biologically relevant processes (Figure 5d;Supplementary Table S19). In the NANOG-centered module, terms passing the adjusted *P <* 0.05 threshold included leukotriene metabolic process (adjusted *P* = 0.0398), regulation of cytokine-mediated signaling pathway (adjusted *P* = 0.0465), regulation of mRNA catabolic process (adjusted *P* = 0.0465), and positive regulation of DNA-templated transcription (adjusted *P* = 0.0465). A related transcriptional term, regulation of transcription by RNA polymerase II, was also among the top-ranked terms but was marginal after adjustment (adjusted *P* = 0.0589), and should therefore be interpreted as suggestive rather than formally significant under a 0.05 adjusted-*P* threshold. Although these enrichment results should be interpreted as module-level functional summaries rather than direct experimental validation, they support the view that BRIDGE-GRN-derived regulatory modules capture coherent transcriptional and signaling-related biological structure.

The biological interpretation analyses show that BRIDGE-GRN learns role-structured latent representations, prioritizes externally supported regulatory edges near the top of its ranking, and organizes high-confidence predictions into functionally interpretable driver-centered modules. These results indicate that the inferred regulatory network is not merely a predictive edge-ranking output, but also provides a structured basis for biological interpretation and downstream hypothesis generation.

## 4 Discussion and Conclusion

### 4.1 Principal findings and methodological implications

This study proposed BRIDGE-GRN, a role-aware graph learning framework for directed gene regulatory network inference from scRNA-seq data. The main finding is that separating shared graph-context encoding from role-specific directional decoding provides a stable and effective strategy for recovering directed TF–target relationships under sparse expression measurements, incomplete prior information, and limited regulatory labels. Across mouse benchmark settings, BRIDGE-GRN achieved consistently strong AUPRC and AUROC performance relative to classical, information-theoretic, motif-guided, dynamical, and recent graph-based baselines. This result is consistent with the broader recognition that scRNA-seq-based GRN inference remains difficult because regulatory positives are sparse, benchmark labels are incomplete, and directionality is hard to recover from expression data alone [11, 12]. Compared with prior graph-based methods such as GENELink, GNNLink, and GCLink, the present framework further emphasizes the explicit decomposition of contextual representation learning and directional role assignment [18–20].

The methodological implication is that directed GRN inference can benefit from treating “gene context”and “regulatory role”as related but distinct modeling objectives. In BRIDGE-GRN, the undirected support graph is used to learn shared gene representations, while directionality is introduced only after projection into TF-role and target-role embedding spaces. This design avoids forcing the graph encoder itself to encode all directional information, which is difficult when priors are incomplete and supervision is limited. The cross-view contrastive objective further regularizes the model by aligning representations learned from identity and edge-perturbed graph views, thereby reducing dependence on a single observed support graph. These design choices complement earlier GRN inference paradigms based on coexpression, information-theoretic dependence, motif-guided regulons, or pseudo-temporal dynamics [14–16], while remaining compatible with recent efforts to formulate GRN inference as a structured prediction problem over genes and regulatory links [18, 20].

### 4.2 Generalization, transferability, and biological interpretation

The benchmark results suggest that BRIDGE-GRN does not rely on a single favorable prior-network condition. Its advantage was observed across Non-Specific, STRING, and Specific prior families, across five mouse cell types, and across both 500- and 1000-gene settings. This is important because prior quality and positive-edge density can strongly affect GRN inference results, and benchmark performance may otherwise reflect properties of a particular reference network rather than a generally useful inference strategy [11, 12]. The transfer analysis further showed that pretrained BRIDGE-GRN representations were most beneficial under low target-domain supervision, supporting the view that regulatory representations learned in one domain can provide useful initialization for adaptation to another domain. This result is consistent with recent single-cell and multi-omic studies emphasizing transferable regulatory structure, atlas-scale external information, and the use of pretrained or representation-learning models in single-cell biology [8, 21–23, 27].

Beyond predictive performance, the biological interpretation analyses showed that BRIDGE-GRN outputs can be examined at multiple levels. The separation between TF-role and target-role embeddings supports the intended role-aware architecture, while the enrichment of top-ranked predictions under curated regulatory support suggests that high-confidence edges are not merely arbitrary ranking artifacts. At the same time, the weak STRING-overlap signal indicates that broad protein-association resources and directed regulatory references capture different forms of biological evidence, and should not be interpreted interchangeably [28]. The recovery of compact driver-centered modules, together with functional enrichment of module targets, suggests that BRIDGE-GRN can provide a hypothesis-generating representation of regulatory programs. This interpretation is aligned with recent work emphasizing that single-cell GRN models should support biological explanation, not only numerical benchmarking [3, 4, 24].

### 4.3 Limitations, future directions, and conclusion

Several limitations should be considered when interpreting these results. First, BRIDGE-GRN still depends on available regulatory labels and prior-network information;therefore, its predictions are constrained by the incompleteness and possible bias of existing references. Second, sampled negative pairs from unlabeled TF–target combinations should not be treated as experimentally confirmed non-regulatory interactions, which is a general challenge in supervised GRN inference. Third, although the support graph was constructed only from training positives to avoid leakage, benchmark references themselves remain incomplete and may favor methods that recover edges similar to the available labels [11, 12]. Fourth, the biological interpretation analyses provide computational evidence of coherence, but they do not constitute direct experimental validation of regulatory causality. This distinction is important because motif support, protein-association overlap, and enrichment analysis can strengthen confidence in candidate modules but cannot replace perturbation-based validation [16, 24, 28].

Future work could extend BRIDGE-GRN in several directions. Integrating perturbation data, chromatin accessibility, enhancer–gene linking, or spatial regulatory information may further improve causal interpretation and context-specific regulatory inference [3, 4, 22, 23]. More extensive cross-species and cross-tissue evaluation would also help determine when pretrained regulatory representations transfer reliably and when domain-specific retraining is required. If BRIDGE-GRN-derived regulatory programs are further extended to disease-risk or outcome-oriented applications, such extensions should be accompanied by careful assessment of data quality, external validity, and predictive uncertainty, as emphasized in broader biomedical prediction and projection studies [29]. Finally, uncertainty calibration and experimentally guided validation could make predicted edges more actionable for downstream biological studies. In conclusion, BRIDGE-GRN provides a compact and interpretable framework for directed GRN inference by combining shared graph-context encoding, role-specific directional decoding, and cross-view regularization. The results support its utility as a robust, transferable, and biologically interpretable approach for inferring directed regulatory structure from single-cell transcriptomic data.

## Author Contributions

H.C. and W.D. contributed equally to this work. H.C. and W.D. jointly conceived the study, designed the methodology, performed the analyses, interpreted the results, prepared the figures and supplementary materials, and wrote and revised the manuscript. Both authors have read and approved the final manuscript.

## Supplementary Information

### 1 Supplementary Note 1. Study design, datasets, and evaluation frame-work

This section summarizes the evaluation framework used for BRIDGE-GRN. Following the model overview, the empirical analyses were organized into four modules: mouse benchmark performance, cross-species low-supervision transfer, architectural dissection, and biological interpretation. These modules evaluated predictive performance, transferability, architectural contributions, and biological interpretability within a unified directed GRN inference framework.

**Table S1.**
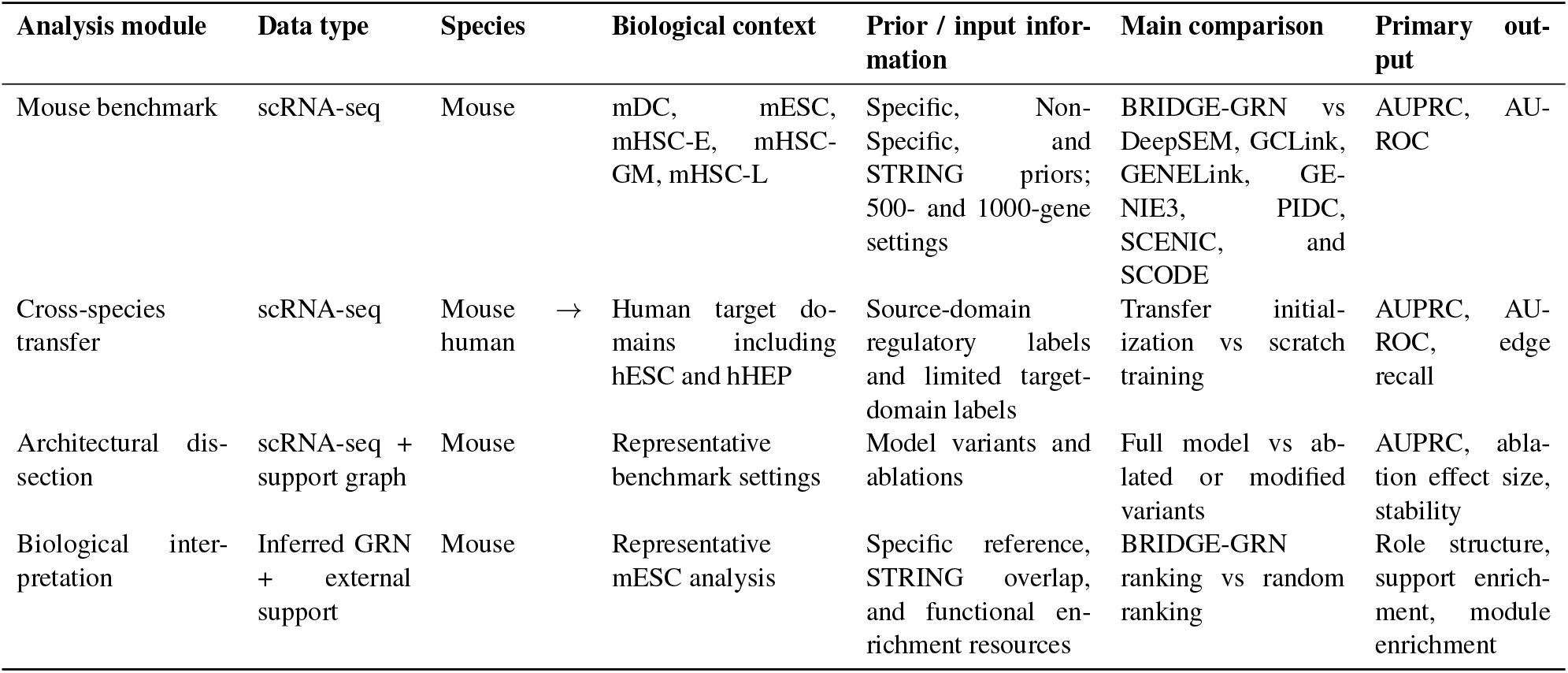
Study design and dataset overview. This table summarizes the empirical analysis modules, data types, species, benchmark or target domains, prior-network information, comparisons, and primary outputs.

**Table S2.**
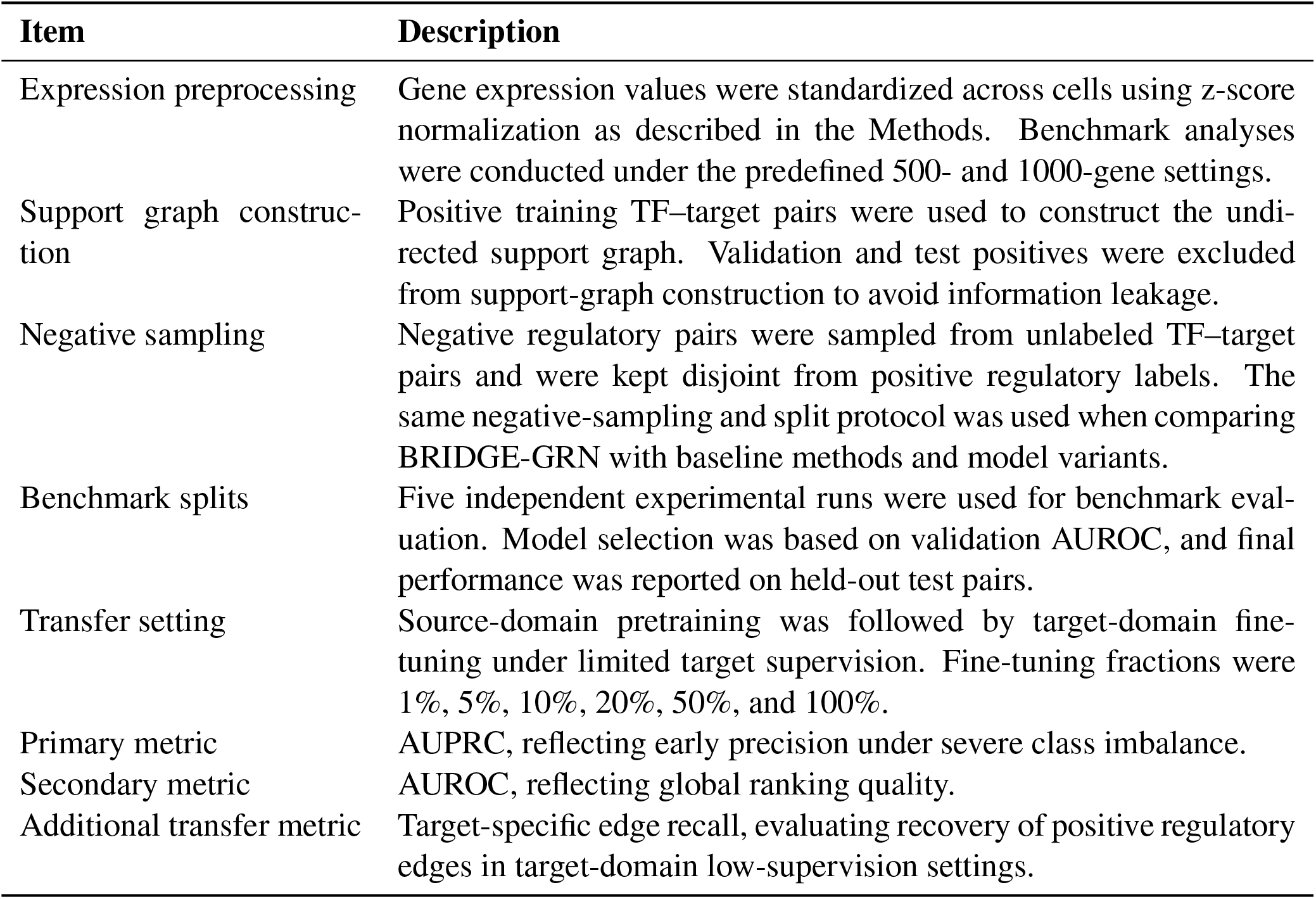
Preprocessing, split strategy, and evaluation metrics. This table summarizes the common preprocessing, graph construction, splitting, transfer-learning, and evaluation settings used in the BRIDGE-GRN analyses.

| Item | Description |
| --- | --- |
| Expression preprocessing | Gene expression values were standardized across cells using z-score normalization as described in the Methods. Benchmark analyses were conducted under the predefined 500- and 1000-gene settings. |
| Support graph construction | Positive training TF–target pairs were used to construct the undirected support graph. Validation and test positives were excluded from support-graph construction to avoid information leakage. |
| Negative sampling | Negative regulatory pairs were sampled from unlabeled TF–target pairs and were kept disjoint from positive regulatory labels. The same negative-sampling and split protocol was used when comparing BRIDGE-GRN with baseline methods and model variants. |
| Benchmark splits | Five independent experimental runs were used for benchmark evaluation. Model selection was based on validation AUROC, and final performance was reported on held-out test pairs. |
| Transfer setting | Source-domain pretraining was followed by target-domain fine-tuning under limited target supervision. Fine-tuning fractions were 1%, 5%, 10%, 20%, 50%, and 100%. |
| Primary metric | AUPRC, reflecting early precision under severe class imbalance. |
| Secondary metric | AUROC, reflecting global ranking quality. |
| Additional transfer metric | Target-specific edge recall, evaluating recovery of positive regulatory edges in target-domain low-supervision settings. |

### 2 Supplementary Note 2. Benchmark performance across mouse GRN settings

This section provides detailed mouse benchmark results supporting the main-text analysis of BRIDGE-GRN generalization across prior-network families, mouse biological contexts, and gene-scale settings.

**Figure S1.**
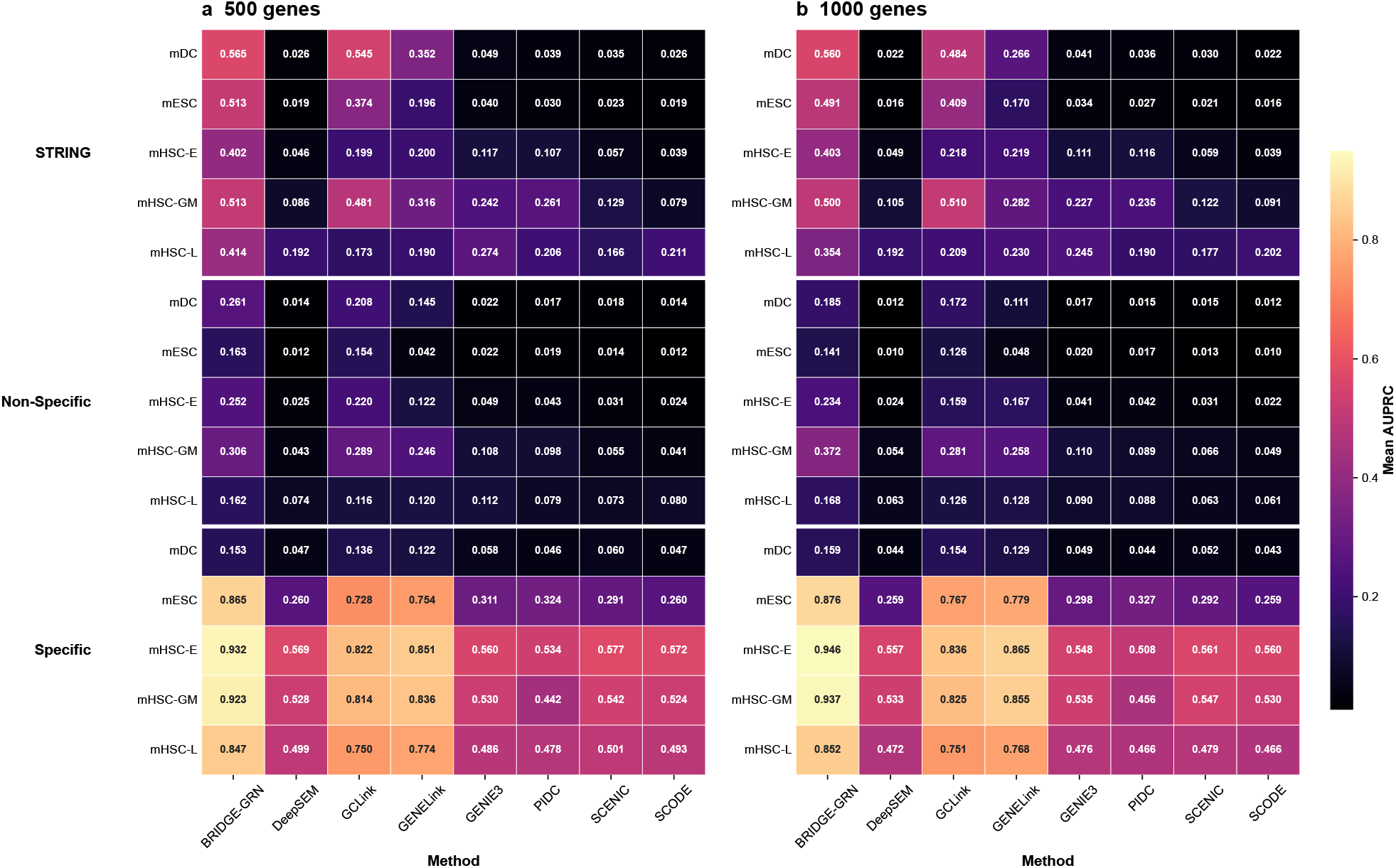
Full AUPRC benchmark performance across mouse GRN settings. Heatmaps show the mean AUPRC across repeated runs for each method under different mouse benchmark configurations. **a**, 500-gene setting. **b**, 1000-gene setting. Rows correspond to combinations of prior-network family and mouse cell type, and columns correspond to GRN inference methods. BRIDGE-GRN is compared with DeepSEM, GCLink, GENELink, GENIE3, PIDC, SCENIC, and SCODE.

**Figure S2.**
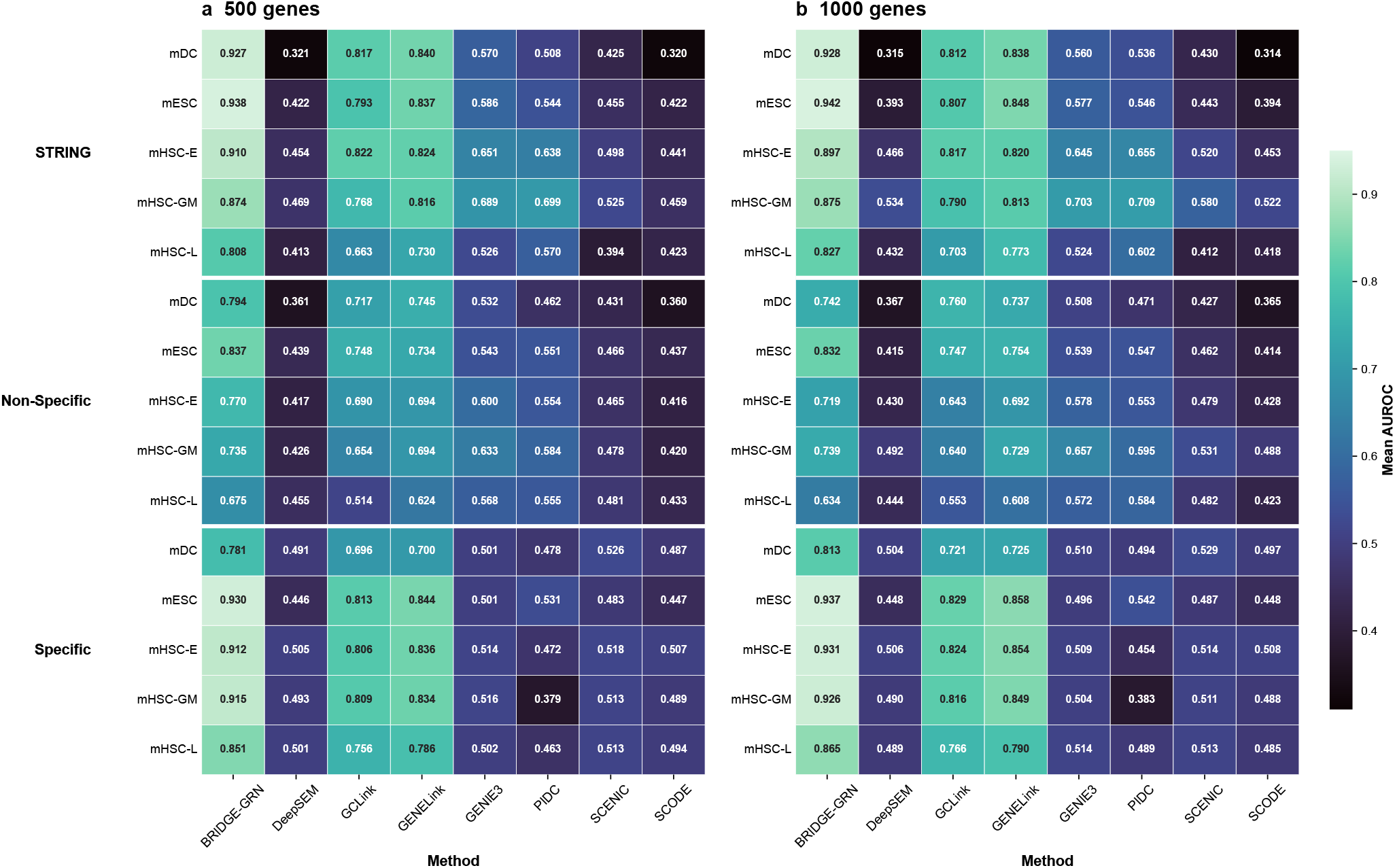
AUROC performance across mouse benchmark configurations. Heatmaps show the mean AUROC across repeated runs for each method under different mouse benchmark configurations. **a**, 500-gene setting. **b**, 1000-gene setting. Rows correspond to combinations of prior-network family and mouse cell type, and columns correspond to GRN inference methods. BRIDGE-GRN is compared with DeepSEM, GCLink, GENELink, GENIE3, PIDC, SCENIC, and SCODE.

**Figure S3.**
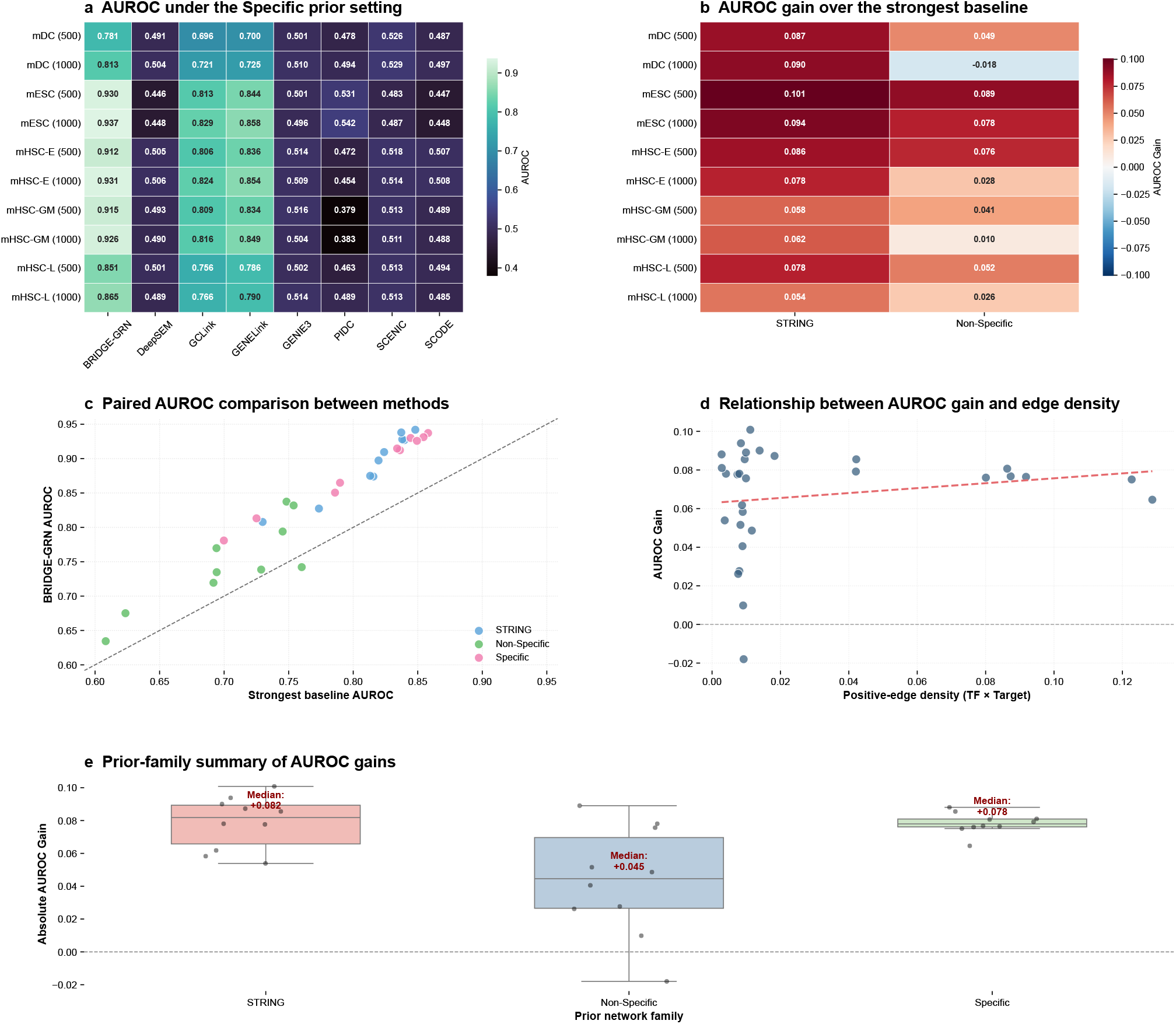
Complementary AUROC consistency analysis across mouse benchmark settings. **a**, AUROC under the Specific prior setting across mouse cell types, gene-scale settings, and GRN inference methods. **b**, AUROC gain of BRIDGE-GRN over the strongest competing baseline under the STRING and Non-Specific prior settings. **c**, Paired AUROC comparison between BRIDGE-GRN and the strongest baseline across benchmark settings. **d**, Relationship between AUROC gain and positive-edge density. **e**, Prior-family summary of AUROC gains across all benchmark settings.

**Table S3.**
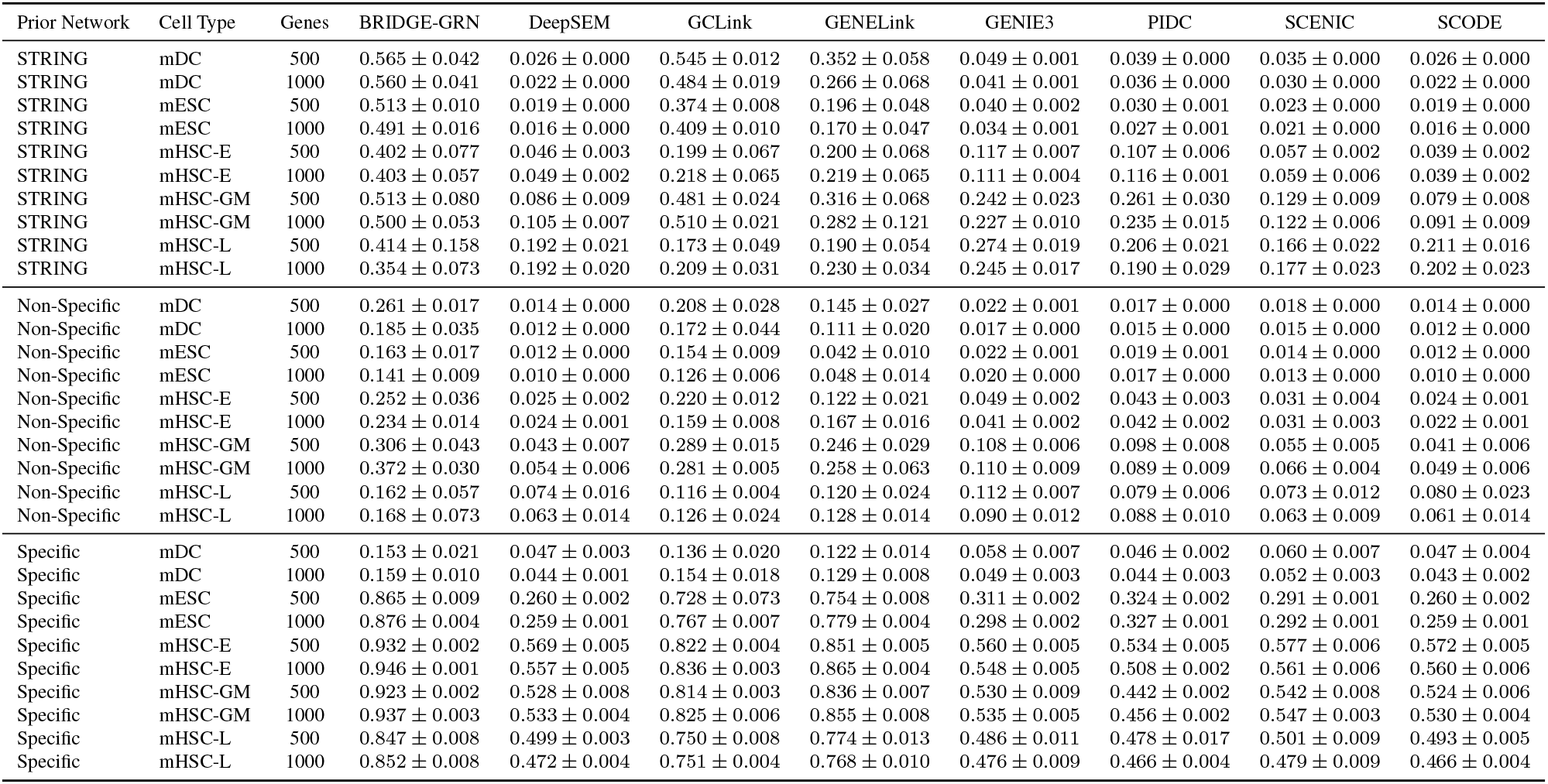
Detailed AUPRC performance across mouse benchmark settings.

**Table S4.**
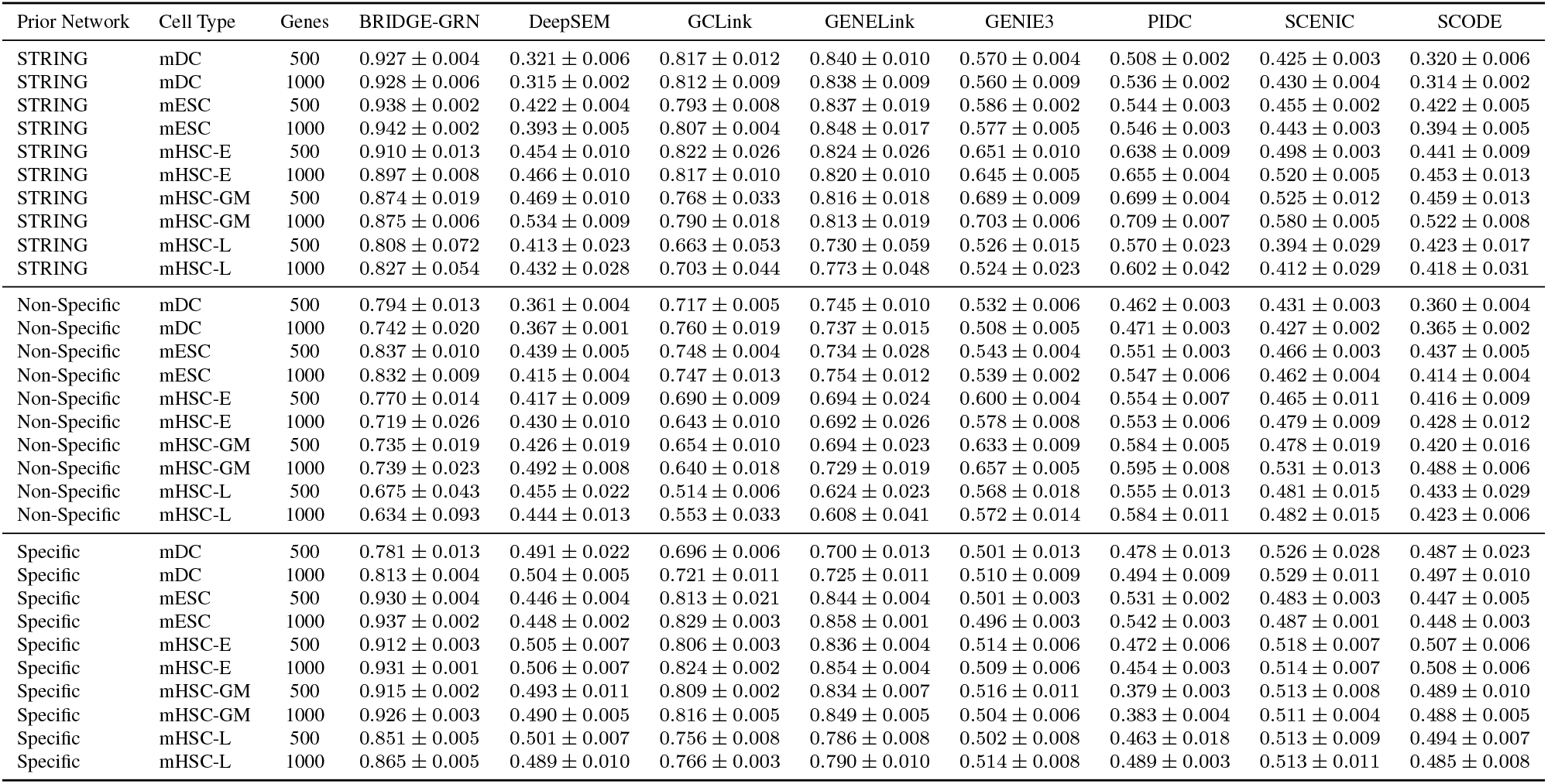
Detailed AUROC performance across mouse benchmark settings.

**Table S5.** AUPRC gains of BRIDGE-GRN over the strongest competing baseline across mouse benchmark settings.

| Prior Network | Cell Type | Genes | Best Baseline AUPRC | BRIDGE-GRN AUPRC | Gain |
| --- | --- | --- | --- | --- | --- |
| STRING | mDC | 500 | GCLink (0.545) | 0.565 | +0.020 |
| STRING | mDC | 1000 | GCLink (0.484) | 0.560 | +0.076 |
| STRING | mESC | 500 | GCLink (0.374) | 0.513 | +0.139 |
| STRING | mESC | 1000 | GCLink (0.409) | 0.491 | +0.082 |
| STRING | mHSC-E | 500 | GENELink (0.200) | 0.402 | +0.202 |
| STRING | mHSC-E | 1000 | GENELink (0.219) | 0.403 | +0.184 |
| STRING | mHSC-GM | 500 | GCLink (0.481) | 0.513 | +0.032 |
| STRING | mHSC-GM | 1000 | GCLink (0.510) | 0.500 | -0.010 |
| STRING | mHSC-L | 500 | GENIE3 (0.274) | 0.414 | +0.140 |
| STRING | mHSC-L | 1000 | GENIE3 (0.245) | 0.354 | +0.108 |
| Non-Specific | mDC | 500 | GCLink (0.208) | 0.261 | +0.052 |
| Non-Specific | mDC | 1000 | GCLink (0.172) | 0.185 | +0.013 |
| Non-Specific | mESC | 500 | GCLink (0.154) | 0.163 | +0.009 |
| Non-Specific | mESC | 1000 | GCLink (0.126) | 0.141 | +0.015 |
| Non-Specific | mHSC-E | 500 | GCLink (0.220) | 0.252 | +0.032 |
| Non-Specific | mHSC-E | 1000 | GENELink (0.167) | 0.234 | +0.067 |
| Non-Specific | mHSC-GM | 500 | GCLink (0.289) | 0.306 | +0.017 |
| Non-Specific | mHSC-GM | 1000 | GCLink (0.281) | 0.372 | +0.091 |
| Non-Specific | mHSC-L | 500 | GENELink (0.120) | 0.162 | +0.042 |
| Non-Specific | mHSC-L | 1000 | GENELink (0.128) | 0.168 | +0.040 |
| Specific | mDC | 500 | GCLink (0.136) | 0.153 | +0.018 |
| Specific | mDC | 1000 | GCLink (0.154) | 0.159 | +0.005 |
| Specific | mESC | 500 | GENELink (0.754) | 0.865 | +0.111 |
| Specific | mESC | 1000 | GENELink (0.779) | 0.876 | +0.097 |
| Specific | mHSC-E | 500 | GENELink (0.851) | 0.932 | +0.081 |
| Specific | mHSC-E | 1000 | GENELink (0.865) | 0.946 | +0.081 |
| Specific | mHSC-GM | 500 | GENELink (0.836) | 0.923 | +0.087 |
| Specific | mHSC-GM | 1000 | GENELink (0.855) | 0.937 | +0.082 |
| Specific | mHSC-L | 500 | GENELink (0.774) | 0.847 | +0.073 |
| Specific | mHSC-L | 1000 | GENELink (0.768) | 0.852 | +0.084 |

**Table S6.** AUROC gains of BRIDGE-GRN over the strongest competing baseline across mouse benchmark settings.

| Prior Network | Cell Type | Genes | Best Baseline AUROC | BRIDGE-GRN AUROC | Gain |
| --- | --- | --- | --- | --- | --- |
| STRING | mDC | 500 | GENELink (0.840) | 0.927 | +0.087 |
| STRING | mDC | 1000 | GENELink (0.838) | 0.928 | +0.090 |
| STRING | mESC | 500 | GENELink (0.837) | 0.938 | +0.101 |
| STRING | mESC | 1000 | GENELink (0.848) | 0.942 | +0.094 |
| STRING | mHSC-E | 500 | GENELink (0.824) | 0.910 | +0.086 |
| STRING | mHSC-E | 1000 | GENELink (0.820) | 0.897 | +0.078 |
| STRING | mHSC-GM | 500 | GENELink (0.816) | 0.874 | +0.058 |
| STRING | mHSC-GM | 1000 | GENELink (0.813) | 0.875 | +0.062 |
| STRING | mHSC-L | 500 | GENELink (0.730) | 0.808 | +0.078 |
| STRING | mHSC-L | 1000 | GENELink (0.773) | 0.827 | +0.054 |
| Non-Specific | mDC | 500 | GENELink (0.745) | 0.794 | +0.049 |
| Non-Specific | mDC | 1000 | GCLink (0.760) | 0.742 | -0.018 |
| Non-Specific | mESC | 500 | GCLink (0.748) | 0.837 | +0.089 |
| Non-Specific | mESC | 1000 | GENELink (0.754) | 0.832 | +0.078 |
| Non-Specific | mHSC-E | 500 | GENELink (0.694) | 0.770 | +0.076 |
| Non-Specific | mHSC-E | 1000 | GENELink (0.692) | 0.719 | +0.028 |
| Non-Specific | mHSC-GM | 500 | GENELink (0.694) | 0.735 | +0.041 |
| Non-Specific | mHSC-GM | 1000 | GENELink (0.729) | 0.739 | +0.010 |
| Non-Specific | mHSC-L | 500 | GENELink (0.624) | 0.675 | +0.052 |
| Non-Specific | mHSC-L | 1000 | GENELink (0.608) | 0.634 | +0.026 |
| Specific | mDC | 500 | GENELink (0.700) | 0.781 | +0.081 |
| Specific | mDC | 1000 | GENELink (0.725) | 0.813 | +0.088 |
| Specific | mESC | 500 | GENELink (0.844) | 0.930 | +0.086 |
| Specific | mESC | 1000 | GENELink (0.858) | 0.937 | +0.079 |
| Specific | mHSC-E | 500 | GENELink (0.836) | 0.912 | +0.076 |
| Specific | mHSC-E | 1000 | GENELink (0.854) | 0.931 | +0.077 |
| Specific | mHSC-GM | 500 | GENELink (0.834) | 0.915 | +0.081 |
| Specific | mHSC-GM | 1000 | GENELink (0.849) | 0.926 | +0.076 |
| Specific | mHSC-L | 500 | GENELink (0.786) | 0.851 | +0.065 |
| Specific | mHSC-L | 1000 | GENELink (0.790) | 0.865 | +0.075 |

**Table S7.** Mouse benchmark network statistics and positive-edge density.

| Prior Type | Cell Type | Genes | Unique Nodes | TFs | Targets | Label Edges | Positive-Edge Density |
| --- | --- | --- | --- | --- | --- | --- | --- |
| STRING | mDC | 500 | 487 | 321 | 821 | 4815 | 0.01827 |
| STRING | mDC | 1000 | 681 | 321 | 1321 | 5898 | 0.01391 |
| STRING | mESC | 500 | 648 | 620 | 1120 | 7762 | 0.01118 |
| STRING | mESC | 1000 | 799 | 620 | 1620 | 8479 | 0.00844 |
| STRING | mHSC-E | 500 | 300 | 204 | 704 | 1371 | 0.00955 |
| STRING | mHSC-E | 1000 | 427 | 204 | 1204 | 1826 | 0.00743 |
| STRING | mHSC-GM | 500 | 206 | 132 | 632 | 748 | 0.00897 |
| STRING | mHSC-GM | 1000 | 357 | 132 | 1132 | 1311 | 0.00877 |
| STRING | mHSC-L | 500 | 74 | 60 | 560 | 137 | 0.00408 |
| STRING | mHSC-L | 1000 | 86 | 60 | 692 | 154 | 0.00371 |
| Non-Specific | mDC | 500 | 643 | 321 | 821 | 3067 | 0.01164 |
| Non-Specific | mDC | 1000 | 980 | 321 | 1321 | 3918 | 0.00924 |
| Non-Specific | mESC | 500 | 896 | 620 | 1120 | 6893 | 0.00993 |
| Non-Specific | mESC | 1000 | 1221 | 620 | 1620 | 8030 | 0.00799 |
| Non-Specific | mHSC-E | 500 | 447 | 204 | 704 | 1425 | 0.00992 |
| Non-Specific | mHSC-E | 1000 | 680 | 204 | 1204 | 1960 | 0.00798 |
| Non-Specific | mHSC-GM | 500 | 301 | 132 | 632 | 743 | 0.00891 |
| Non-Specific | mHSC-GM | 1000 | 532 | 132 | 1132 | 1358 | 0.00909 |
| Non-Specific | mHSC-L | 500 | 168 | 60 | 560 | 279 | 0.00830 |
| Non-Specific | mHSC-L | 1000 | 198 | 60 | 692 | 317 | 0.00763 |
| Specific | mDC | 500 | 448 | 323 | 821 | 756 | 0.00285 |
| Specific | mDC | 1000 | 690 | 323 | 1321 | 1193 | 0.00280 |
| Specific | mESC | 500 | 977 | 627 | 1120 | 29613 | 0.04217 |
| Specific | mESC | 1000 | 1385 | 628 | 1620 | 42795 | 0.04206 |
| Specific | mHSC-E | 500 | 691 | 205 | 704 | 11557 | 0.08008 |
| Specific | mHSC-E | 1000 | 1177 | 209 | 1204 | 21975 | 0.08733 |
| Specific | mHSC-GM | 500 | 618 | 135 | 632 | 7364 | 0.08631 |
| Specific | mHSC-GM | 1000 | 1089 | 136 | 1132 | 14135 | 0.09181 |
| Specific | mHSC-L | 500 | 525 | 61 | 560 | 4398 | 0.12875 |
| Specific | mHSC-L | 1000 | 640 | 61 | 692 | 5180 | 0.12271 |

**Table S8.** Global transfer-learning summary across fine-tuning fractions.

| Fraction | Scratch AUPRC | Transfer AUPRC | AUPRC Gain | Scratch AUROC | Transfer AUROC | AUROC Gain | Runs |
| --- | --- | --- | --- | --- | --- | --- | --- |
| 1% | 0.426 ± 0.231 | 0.515 ± 0.225 | 0.089 ± 0.080 | 0.548 ± 0.135 | 0.719 ± 0.073 | 0.171 ± 0.149 | 35 |
| 5% | 0.488 ± 0.248 | 0.596 ± 0.251 | 0.108 ± 0.060 | 0.561 ± 0.136 | 0.793 ± 0.046 | 0.232 ± 0.135 | 35 |
| 10% | 0.569 ± 0.280 | 0.627 ± 0.263 | 0.058 ± 0.050 | 0.687 ± 0.110 | 0.806 ± 0.055 | 0.119 ± 0.076 | 35 |
| 20% | 0.648 ± 0.280 | 0.675 ± 0.265 | 0.027 ± 0.026 | 0.793 ± 0.090 | 0.843 ± 0.048 | 0.050 ± 0.047 | 35 |
| 50% | 0.714 ± 0.268 | 0.713 ± 0.274 | -0.001 ± 0.016 | 0.865 ± 0.056 | 0.869 ± 0.055 | 0.004 ± 0.018 | 35 |
| 100% | 0.740 ± 0.265 | 0.743 ± 0.265 | 0.004 ± 0.014 | 0.888 ± 0.047 | 0.889 ± 0.049 | 0.002 ± 0.012 | 35 |

**Table S9.** Human target-domain transfer summary for hESC and hHEP.

| Cell Type | Fraction | Scratch AUPRC | Transfer AUPRC | AUPRC Gain | Scratch AUROC | Transfer AUROC | AUROC Gain | Runs |
| --- | --- | --- | --- | --- | --- | --- | --- | --- |
| hESC | 1% | 0.201 ± 0.041 | 0.440 ± 0.015 | 0.239 ± 0.038 | 0.431 ± 0.084 | 0.803 ± 0.007 | 0.372 ± 0.081 | 5 |
| hESC | 5% | 0.184 ± 0.010 | 0.382 ± 0.011 | 0.198 ± 0.014 | 0.342 ± 0.009 | 0.794 ± 0.008 | 0.451 ± 0.015 | 5 |
| hESC | 10% | 0.252 ± 0.027 | 0.399 ± 0.019 | 0.147 ± 0.041 | 0.596 ± 0.043 | 0.798 ± 0.005 | 0.202 ± 0.045 | 5 |
| hESC | 20% | 0.424 ± 0.030 | 0.472 ± 0.022 | 0.048 ± 0.043 | 0.782 ± 0.011 | 0.820 ± 0.013 | 0.039 ± 0.020 | 5 |
| hESC | 50% | 0.547 ± 0.009 | 0.515 ± 0.010 | -0.032 ± 0.012 | 0.845 ± 0.007 | 0.834 ± 0.006 | -0.011 ± 0.008 | 5 |
| hESC | 100% | 0.604 ± 0.010 | 0.587 ± 0.011 | -0.018 ± 0.007 | 0.876 ± 0.004 | 0.867 ± 0.004 | -0.009 ± 0.006 | 5 |
| hHEP | 1% | 0.400 ± 0.018 | 0.489 ± 0.024 | 0.089 ± 0.028 | 0.472 ± 0.052 | 0.696 ± 0.038 | 0.224 ± 0.054 | 5 |
| hHEP | 5% | 0.489 ± 0.006 | 0.620 ± 0.014 | 0.131 ± 0.011 | 0.480 ± 0.011 | 0.771 ± 0.011 | 0.291 ± 0.011 | 5 |
| hHEP | 10% | 0.603 ± 0.012 | 0.684 ± 0.010 | 0.081 ± 0.015 | 0.627 ± 0.009 | 0.801 ± 0.007 | 0.174 ± 0.011 | 5 |
| hHEP | 20% | 0.703 ± 0.011 | 0.729 ± 0.013 | 0.026 ± 0.009 | 0.797 ± 0.011 | 0.833 ± 0.008 | 0.036 ± 0.008 | 5 |
| hHEP | 50% | 0.795 ± 0.004 | 0.800 ± 0.005 | 0.005 ± 0.005 | 0.876 ± 0.005 | 0.880 ± 0.003 | 0.004 ± 0.006 | 5 |
| hHEP | 100% | 0.833 ± 0.003 | 0.829 ± 0.007 | -0.004 ± 0.006 | 0.898 ± 0.003 | 0.896 ± 0.004 | -0.003 ± 0.005 | 5 |

**Table S10.** Cell-type-specific transfer-learning summary across fine-tuning fractions.

| Cell Type | Fraction | Scratch AUPRC | Transfer AUPRC | AUPRC Gain | Scratch AUROC | Transfer AUROC | AUROC Gain | Runs |
| --- | --- | --- | --- | --- | --- | --- | --- | --- |
| hESC | 1% | 0.201 ± 0.041 | 0.440 ± 0.015 | 0.239 ± 0.038 | 0.431 ± 0.084 | 0.803 ± 0.007 | 0.372 ± 0.081 | 5 |
| hESC | 5% | 0.184 ± 0.010 | 0.382 ± 0.011 | 0.198 ± 0.014 | 0.342 ± 0.009 | 0.794 ± 0.008 | 0.451 ± 0.015 | 5 |
| hESC | 10% | 0.252 ± 0.027 | 0.399 ± 0.019 | 0.147 ± 0.041 | 0.596 ± 0.043 | 0.798 ± 0.005 | 0.202 ± 0.045 | 5 |
| hESC | 20% | 0.424 ± 0.030 | 0.472 ± 0.022 | 0.048 ± 0.043 | 0.782 ± 0.011 | 0.820 ± 0.013 | 0.039 ± 0.020 | 5 |
| hESC | 50% | 0.547 ± 0.009 | 0.515 ± 0.010 | -0.032 ± 0.012 | 0.845 ± 0.007 | 0.834 ± 0.006 | -0.011 ± 0.008 | 5 |
| hESC | 100% | 0.604 ± 0.010 | 0.587 ± 0.011 | -0.018 ± 0.007 | 0.876 ± 0.004 | 0.867 ± 0.004 | -0.009 ± 0.006 | 5 |
| hHEP | 1% | 0.400 ± 0.018 | 0.489 ± 0.024 | 0.089 ± 0.028 | 0.472 ± 0.052 | 0.696 ± 0.038 | 0.224 ± 0.054 | 5 |
| hHEP | 5% | 0.489 ± 0.006 | 0.620 ± 0.014 | 0.131 ± 0.011 | 0.480 ± 0.011 | 0.771 ± 0.011 | 0.291 ± 0.011 | 5 |
| hHEP | 10% | 0.603 ± 0.012 | 0.684 ± 0.010 | 0.081 ± 0.015 | 0.627 ± 0.009 | 0.801 ± 0.007 | 0.174 ± 0.011 | 5 |
| hHEP | 20% | 0.703 ± 0.011 | 0.729 ± 0.013 | 0.026 ± 0.009 | 0.797 ± 0.011 | 0.833 ± 0.008 | 0.036 ± 0.008 | 5 |
| hHEP | 50% | 0.795 ± 0.004 | 0.800 ± 0.005 | 0.005 ± 0.005 | 0.876 ± 0.005 | 0.880 ± 0.003 | 0.004 ± 0.006 | 5 |
| hHEP | 100% | 0.833 ± 0.003 | 0.829 ± 0.007 | -0.004 ± 0.006 | 0.898 ± 0.003 | 0.896 ± 0.004 | -0.003 ± 0.005 | 5 |
| mDC | 1% | 0.061 ± 0.004 | 0.064 ± 0.011 | 0.002 ± 0.012 | 0.600 ± 0.043 | 0.605 ± 0.073 | 0.005 ± 0.088 | 5 |
| mDC | 5% | 0.076 ± 0.010 | 0.105 ± 0.015 | 0.028 ± 0.007 | 0.695 ± 0.018 | 0.749 ± 0.013 | 0.054 ± 0.020 | 5 |
| mDC | 10% | 0.070 ± 0.008 | 0.103 ± 0.010 | 0.033 ± 0.016 | 0.681 ± 0.035 | 0.741 ± 0.017 | 0.060 ± 0.042 | 5 |
| mDC | 20% | 0.082 ± 0.013 | 0.129 ± 0.006 | 0.047 ± 0.013 | 0.671 ± 0.042 | 0.782 ± 0.004 | 0.111 ± 0.042 | 5 |
| mDC | 50% | 0.136 ± 0.014 | 0.135 ± 0.010 | -0.001 ± 0.010 | 0.780 ± 0.006 | 0.772 ± 0.019 | -0.008 ± 0.023 | 5 |
| mDC | 100% | 0.153 ± 0.009 | 0.165 ± 0.008 | 0.012 ± 0.010 | 0.801 ± 0.009 | 0.795 ± 0.007 | -0.007 ± 0.006 | 5 |
| mESC | 1% | 0.366 ± 0.008 | 0.474 ± 0.011 | 0.108 ± 0.013 | 0.386 ± 0.011 | 0.695 ± 0.018 | 0.309 ± 0.017 | 5 |
| mESC | 5% | 0.638 ± 0.011 | 0.714 ± 0.011 | 0.075 ± 0.009 | 0.684 ± 0.014 | 0.835 ± 0.006 | 0.150 ± 0.011 | 5 |
| mESC | 10% | 0.779 ± 0.006 | 0.779 ± 0.009 | 0.000 ± 0.005 | 0.865 ± 0.006 | 0.872 ± 0.007 | 0.007 ± 0.005 | 5 |
| mESC | 20% | 0.825 ± 0.006 | 0.834 ± 0.004 | 0.009 ± 0.004 | 0.908 ± 0.003 | 0.911 ± 0.003 | 0.002 ± 0.002 | 5 |
| mESC | 50% | 0.852 ± 0.004 | 0.868 ± 0.003 | 0.017 ± 0.006 | 0.925 ± 0.002 | 0.933 ± 0.001 | 0.008 ± 0.003 | 5 |
| mESC | 100% | 0.863 ± 0.004 | 0.883 ± 0.003 | 0.019 ± 0.005 | 0.932 ± 0.003 | 0.940 ± 0.001 | 0.008 ± 0.003 | 5 |
| mHSC-E | 1% | 0.712 ± 0.036 | 0.762 ± 0.033 | 0.050 ± 0.023 | 0.691 ± 0.082 | 0.763 ± 0.035 | 0.073 ± 0.070 | 5 |
| mHSC-E | 5% | 0.761 ± 0.043 | 0.850 ± 0.015 | 0.089 ± 0.047 | 0.630 ± 0.100 | 0.839 ± 0.017 | 0.209 ± 0.103 | 5 |
| mHSC-E | 10% | 0.854 ± 0.015 | 0.880 ± 0.005 | 0.026 ± 0.014 | 0.781 ± 0.034 | 0.865 ± 0.009 | 0.084 ± 0.033 | 5 |
| mHSC-E | 20% | 0.913 ± 0.002 | 0.909 ± 0.003 | -0.004 ± 0.003 | 0.887 ± 0.005 | 0.891 ± 0.002 | 0.004 ± 0.005 | 5 |
| mHSC-E | 50% | 0.936 ± 0.002 | 0.934 ± 0.004 | -0.002 ± 0.005 | 0.920 ± 0.003 | 0.920 ± 0.003 | -0.001 ± 0.006 | 5 |
| mHSC-E | 100% | 0.945 ± 0.004 | 0.947 ± 0.004 | 0.003 ± 0.006 | 0.930 ± 0.004 | 0.933 ± 0.003 | 0.003 ± 0.006 | 5 |
| mHSC-GM | 1% | 0.680 ± 0.070 | 0.753 ± 0.030 | 0.073 ± 0.069 | 0.664 ± 0.131 | 0.780 ± 0.033 | 0.116 ± 0.138 | 5 |
| mHSC-GM | 5% | 0.706 ± 0.041 | 0.834 ± 0.030 | 0.128 ± 0.055 | 0.598 ± 0.099 | 0.835 ± 0.031 | 0.237 ± 0.109 | 5 |
| mHSC-GM | 10% | 0.802 ± 0.033 | 0.845 ± 0.008 | 0.043 ± 0.029 | 0.714 ± 0.075 | 0.837 ± 0.008 | 0.123 ± 0.070 | 5 |
| mHSC-GM | 20% | 0.876 ± 0.015 | 0.891 ± 0.003 | 0.015 ± 0.013 | 0.830 ± 0.032 | 0.878 ± 0.003 | 0.049 ± 0.031 | 5 |
| mHSC-GM | 50% | 0.924 ± 0.005 | 0.923 ± 0.005 | -0.001 ± 0.004 | 0.911 ± 0.005 | 0.912 ± 0.003 | 0.001 ± 0.005 | 5 |
| mHSC-GM | 100% | 0.939 ± 0.002 | 0.939 ± 0.004 | 0.000 ± 0.004 | 0.929 ± 0.001 | 0.928 ± 0.005 | -0.001 ± 0.005 | 5 |
| mHSC-L | 1% | 0.561 ± 0.071 | 0.626 ± 0.040 | 0.065 ± 0.076 | 0.594 ± 0.118 | 0.691 ± 0.036 | 0.097 ± 0.114 | 5 |
| mHSC-L | 5% | 0.561 ± 0.054 | 0.665 ± 0.012 | 0.104 ± 0.062 | 0.493 ± 0.121 | 0.726 ± 0.018 | 0.233 ± 0.135 | 5 |
| mHSC-L | 10% | 0.621 ± 0.017 | 0.697 ± 0.015 | 0.077 ± 0.015 | 0.541 ± 0.017 | 0.726 ± 0.018 | 0.185 ± 0.017 | 5 |
| mHSC-L | 20% | 0.715 ± 0.009 | 0.763 ± 0.008 | 0.047 ± 0.009 | 0.676 ± 0.008 | 0.787 ± 0.010 | 0.111 ± 0.013 | 5 |
| mHSC-L | 50% | 0.807 ± 0.008 | 0.817 ± 0.004 | 0.010 ± 0.012 | 0.798 ± 0.011 | 0.833 ± 0.004 | 0.035 ± 0.014 | 5 |
| mHSC-L | 100% | 0.841 ± 0.010 | 0.854 ± 0.006 | 0.013 ± 0.015 | 0.849 ± 0.015 | 0.869 ± 0.004 | 0.019 ± 0.017 | 5 |

**Table S11.** Target-specific edge recovery summary under low-shot transfer learning.

| Fraction | Method | Target-specific recall | Recovered positive edges | AUPRC | AUROC | Runs |
| --- | --- | --- | --- | --- | --- | --- |
| 1% | Scratch | 0.134 ± 0.106 | 220.4 ± 173.5 | 0.147 ± 0.042 | 0.301 ± 0.070 | 5 |
| 1% | Transfer | 0.419 ± 0.018 | 688.8 ± 30.3 | 0.374 ± 0.014 | 0.779 ± 0.024 | 5 |
| 5% | Scratch | 0.194 ± 0.006 | 318.6 ± 10.5 | 0.221 ± 0.033 | 0.551 ± 0.110 | 5 |
| 5% | Transfer | 0.409 ± 0.031 | 672.4 ± 50.6 | 0.387 ± 0.012 | 0.800 ± 0.006 | 5 |
| 10% | Scratch | 0.383 ± 0.019 | 629.0 ± 31.8 | 0.367 ± 0.017 | 0.782 ± 0.013 | 5 |
| 10% | Transfer | 0.421 ± 0.014 | 692.0 ± 23.0 | 0.418 ± 0.009 | 0.806 ± 0.010 | 5 |

**Table S12.** Latent-space alignment summary statistics for the representative transfer analysis.

| State | Cell Type | Sample | Fraction | Centroid distance | Mean TF norm | Mean target norm | AUPRC | AUROC |
| --- | --- | --- | --- | --- | --- | --- | --- | --- |
| Scratch | hESC | sample1 | 0.050 | 15.878 | 8.746 | 13.163 | 0.212 | 0.563 |
| Transfer | hESC | sample1 | 0.050 | 30.475 | 24.078 | 19.007 | 0.381 | 0.789 |

### 3 Supplementary Note 3. Cross-species transfer and low-supervision generalization

This section provides extended analyses for the mouse-to-human transfer setting and the broader low-supervision transfer experiments. These results support the conclusion that pretrained BRIDGE-GRN representations improve target-domain adaptation when target regulatory labels are scarce.

**Figure S4.**
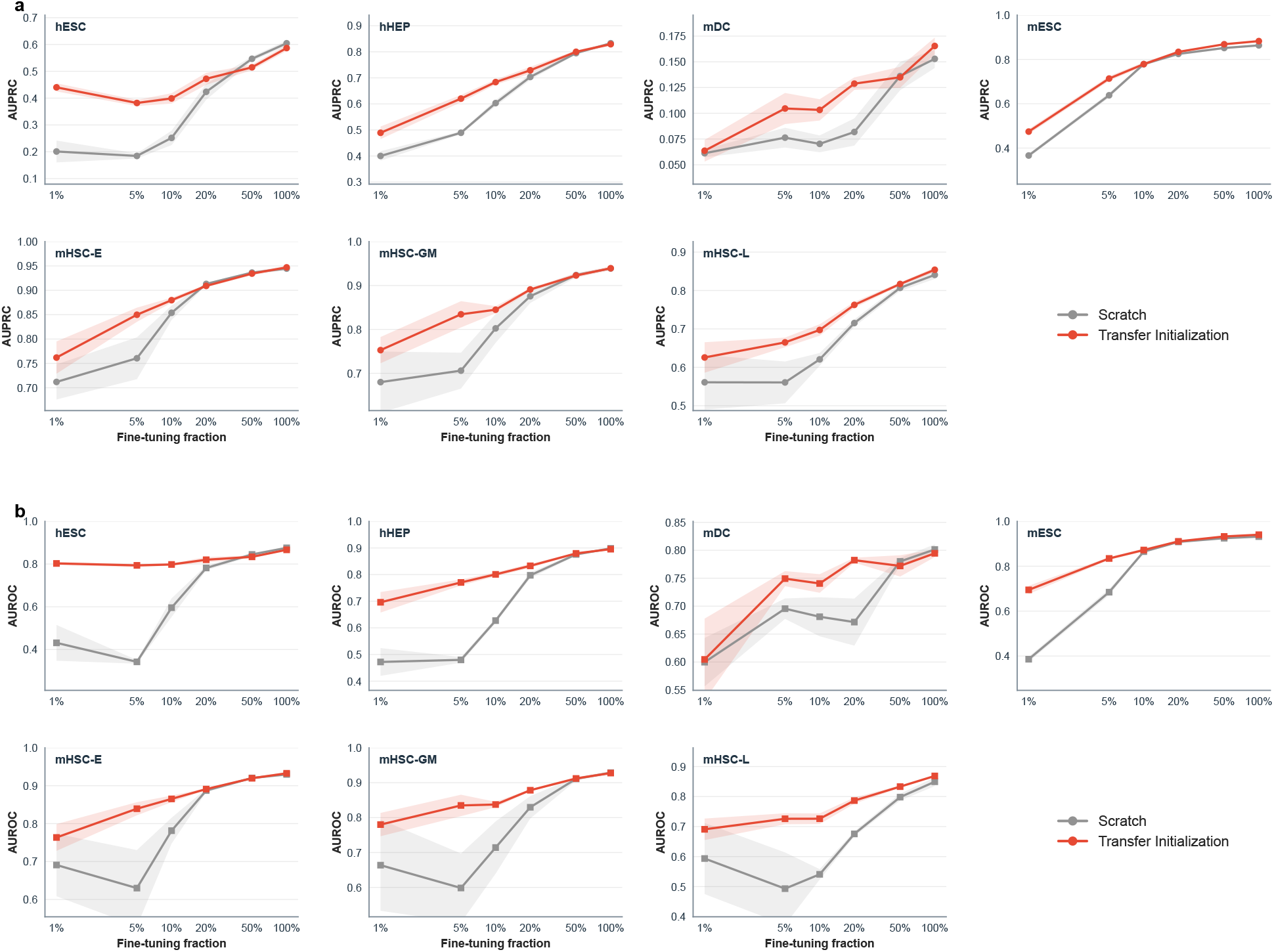
Cell-type-specific transfer-learning performance curves across fine-tuning fractions. **a**, AUPRC curves for each target cell type across fine-tuning fractions from 1%to 100%, comparing scratch training and transfer initialization. **b**, AUROC curves for the same target cell types and fine-tuning fractions. Transfer initialization provides the largest benefit in the low-supervision regime, with gains narrowing as target-domain supervision increases.

### 4 Supplementary Note 4. Architectural ablation and robustness analyses

This section provides detailed ablation and robustness analyses supporting the conclusion that BRIDGE-GRN performance is mainly driven by role-specific directional factorization and cross-view regularization.

### 5 Supplementary Note 5. Biological interpretation and regulatory module analysis

This section provides extended analyses of BRIDGE-GRN latent representations, external support of top-ranked predictions, driver-centered regulatory modules, and functional enrichment.

**Figure S5.**
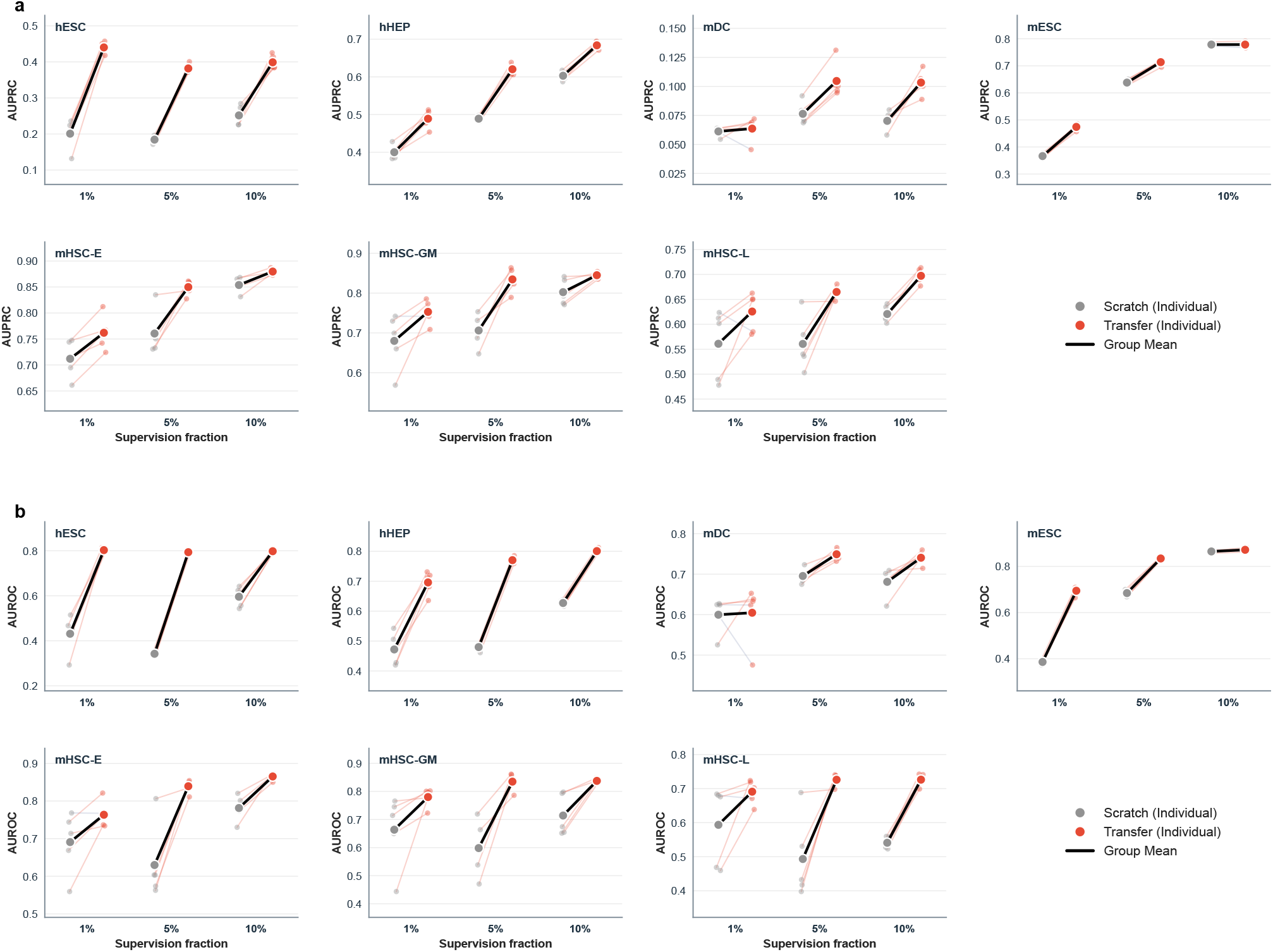
Low-shot paired comparisons under transfer initialization. **a**, Sample-wise paired AUPRC comparisons at 1%, 5%, and 10%target supervision. **b**, Corresponding paired AUROC comparisons. Thin lines connect matched scratch and transfer runs, and bold markers summarize group means.

**Table S13.** Complete ablation numerical results. Results are reported as mean and standard deviation across five repeated runs in the representative Specific–mESC–1000-gene setting. ! values are computed as variant minus full BRIDGE-GRN;therefore, negative values indicate lower performance than the full model.

| Variant | AUPRC mean | AUPRC SD | ΔAUPRC vs full | AUROC mean | AUROC SD | ΔAUROC vs full |
| --- | --- | --- | --- | --- | --- | --- |
| Full BRIDGE-GRN | 0.876 | 0.004 | Reference | 0.937 | 0.002 | Reference |
| Without contrastive regularization | 0.857 | 0.004 | -0.019 | 0.928 | 0.002 | -0.009 |
| Single shared tower | 0.844 | 0.007 | -0.032 | 0.919 | 0.004 | -0.018 |
| Identity-view only | 0.873 | 0.003 | -0.004 | 0.934 | 0.004 | -0.004 |
| Mean fusion instead of concatenation | 0.869 | 0.006 | -0.007 | 0.934 | 0.004 | -0.003 |
| Smaller embedding dimension | 0.868 | 0.006 | -0.008 | 0.934 | 0.002 | -0.003 |
| Cosine decoder | 0.828 | 0.003 | -0.049 | 0.895 | 0.002 | -0.042 |
| Bilinear decoder | 0.872 | 0.006 | -0.004 | 0.930 | 0.001 | -0.007 |

**Figure S6.**
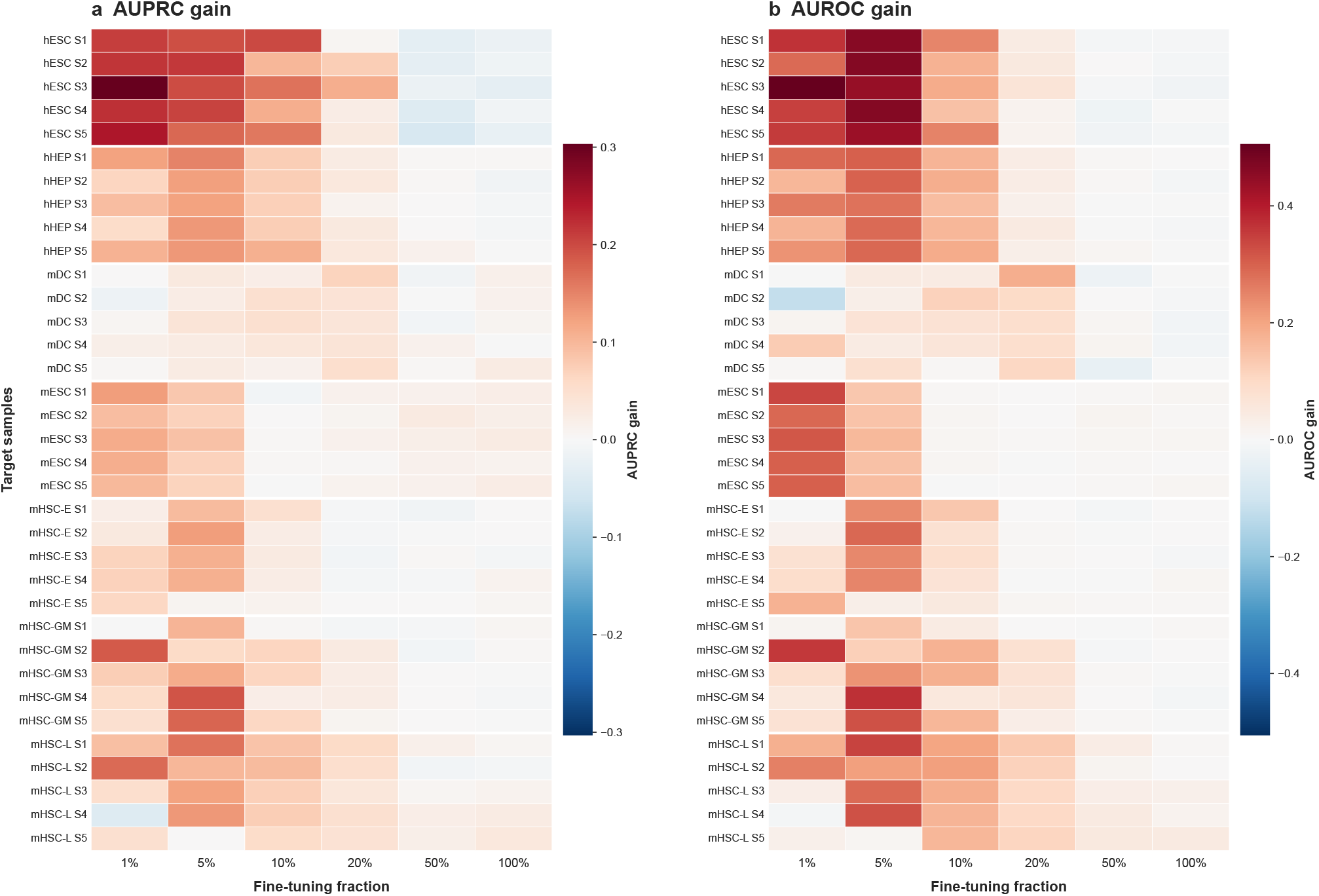
Transfer-gain heatmaps across target-domain samples and fine-tuning fractions. **a**, AUPRC gain, defined as Transfer −Scratch, across target-domain samples and fine-tuning fractions. **b**, AUROC gain, defined as Transfer −Scratch, across the same settings. Positive values indicate improved performance after transfer initialization.

**Figure S7.**
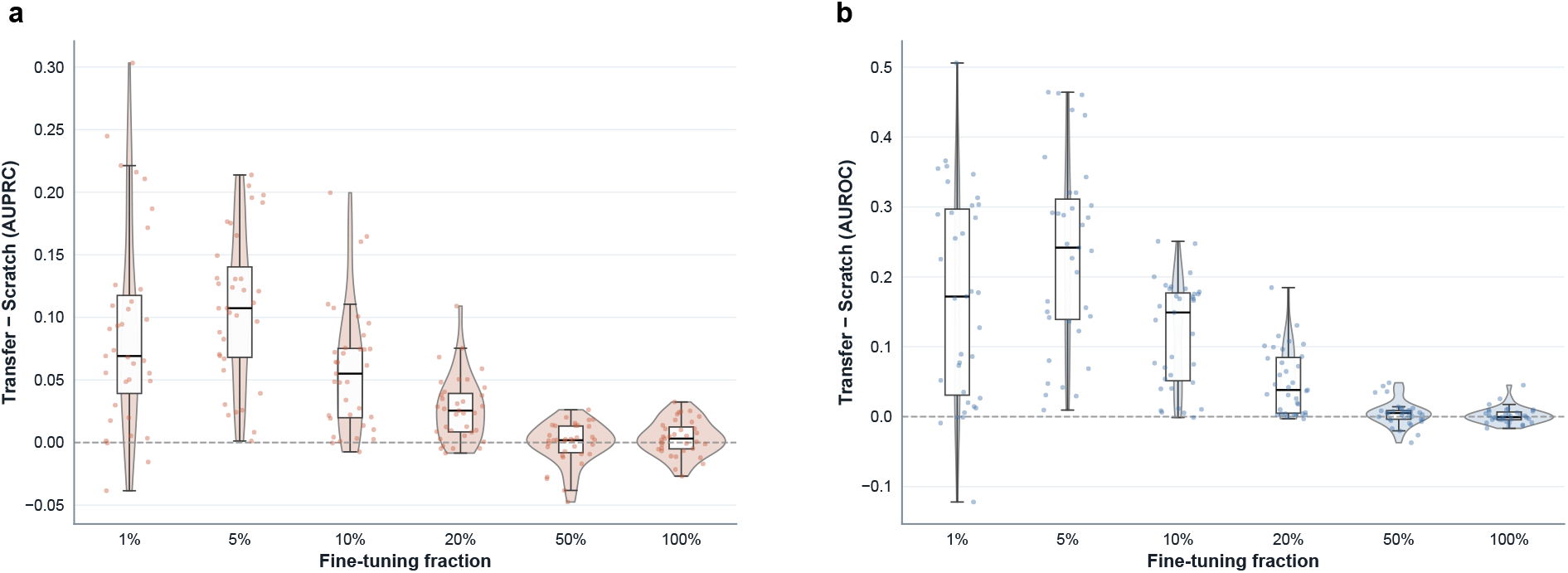
Distribution of transfer gains across fine-tuning fractions. **a**, Distribution of AUPRC gain, defined as Transfer −Scratch, across target-domain runs at each fine-tuning fraction. **b**, Distribution of AUROC gain across the same settings. Violin plots, boxplots, and jittered points jointly show that transfer gains are strongest in the low-supervision regime and narrow as target-domain supervision increases.

**Figure S8.**
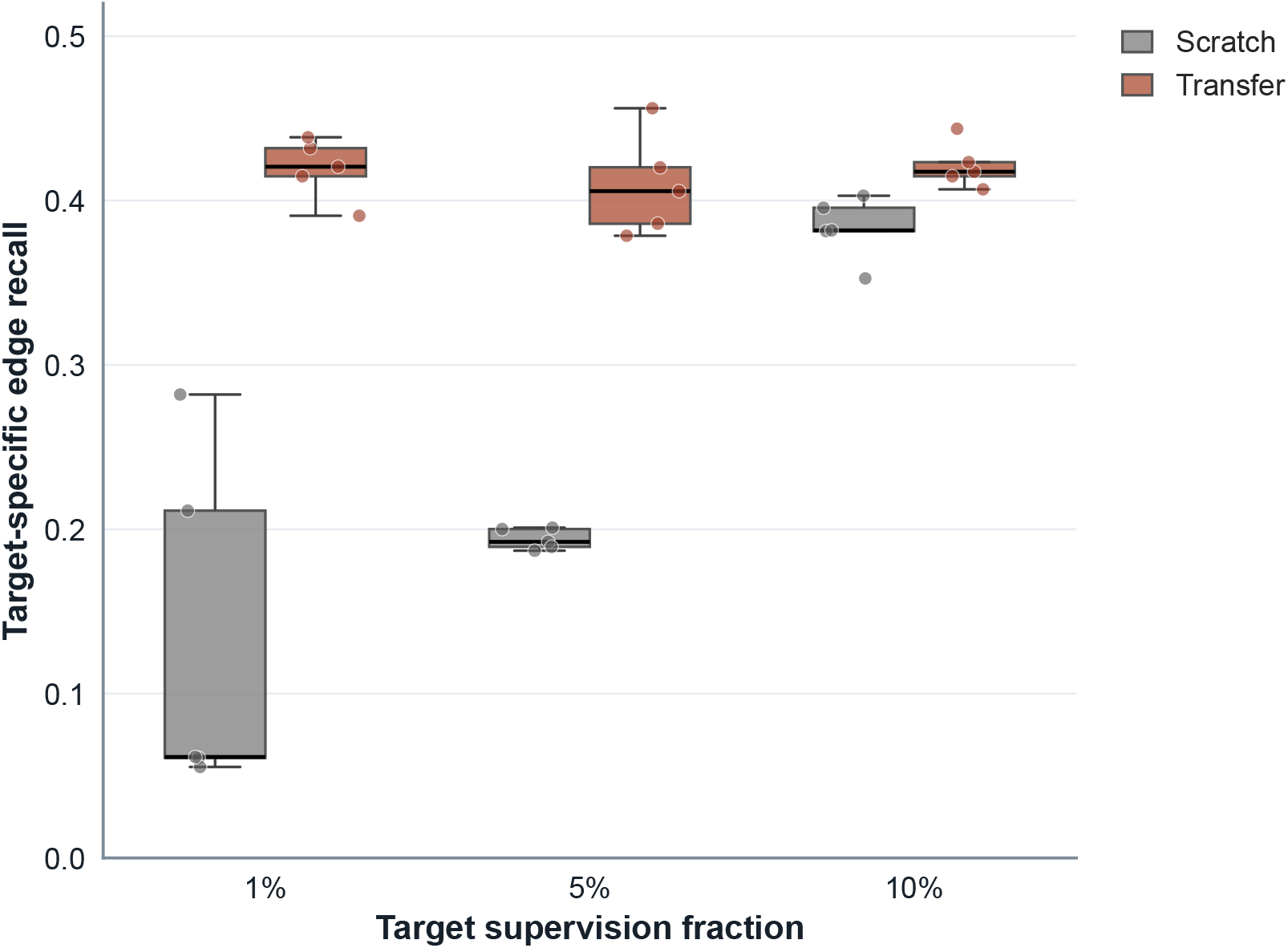
Target-specific edge recovery under low-shot transfer learning. Target-specific edge recall is shown for scratch training and transfer initialization at 1%, 5%, and 10%target supervision. Boxplots summarize the distribution across repeated runs, and jittered points show individual runs.

**Figure S9.**
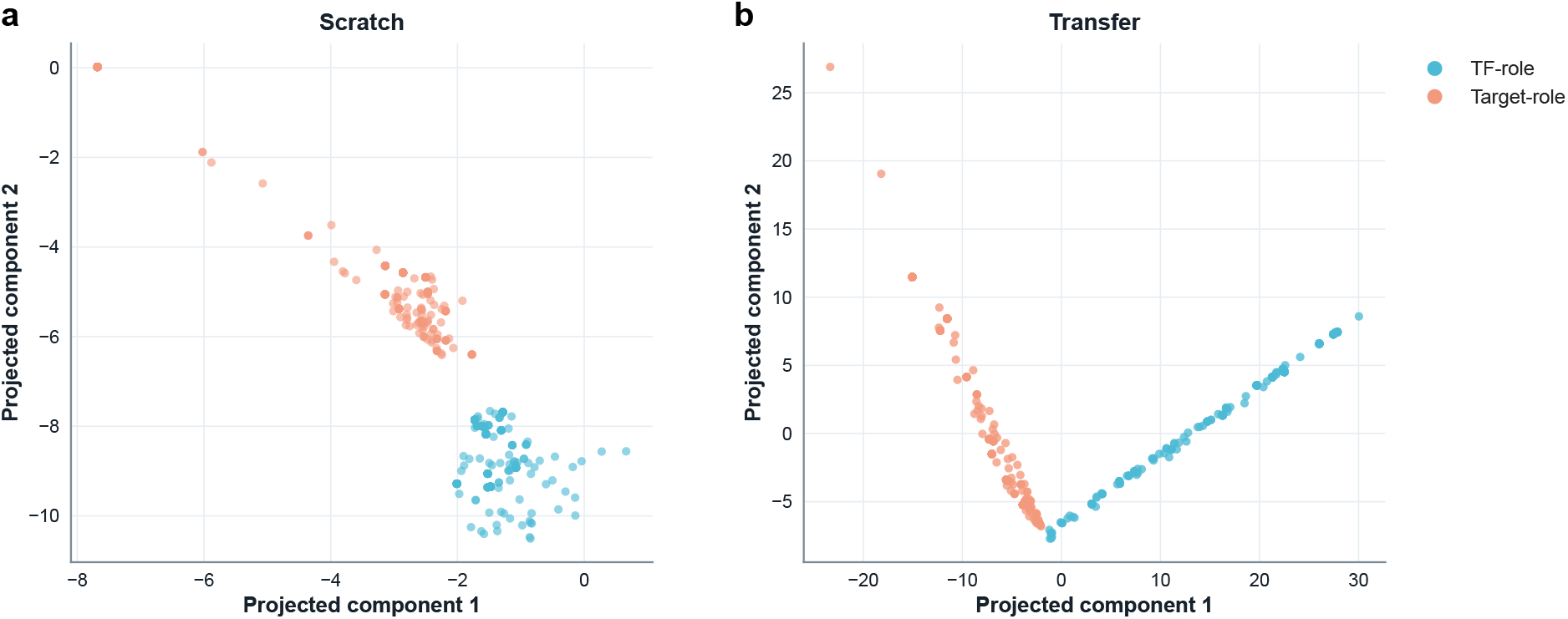
Representative latent-space projections under scratch and transfer initialization. **a**, Latent projection learned from scratch at 5%target-domain supervision. **b**, Corresponding latent projection obtained with transfer initialization. TF-role and target-role node embeddings are shown separately.

**Figure S10.**
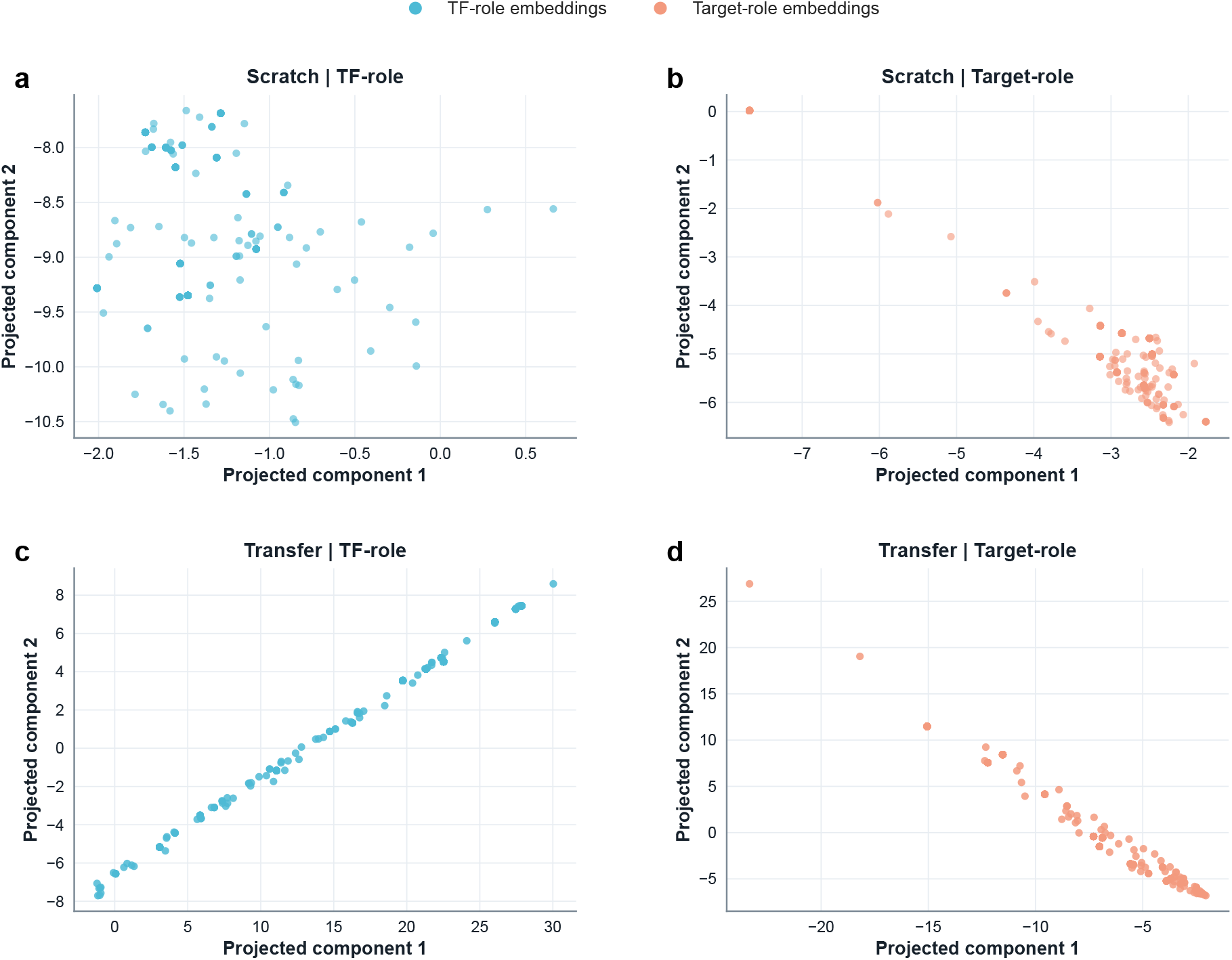
Additional role-specific latent projections in the transfer analysis. **a**, Scratch TF-role embeddings. **b**, Scratch target-role embeddings. **c**, Transfer TF-role embeddings. **d**, Transfer target-role embeddings. These projections provide a role-specific decomposition of the latent-space organization shown in Supplementary Figure S9.

**Figure S11.**
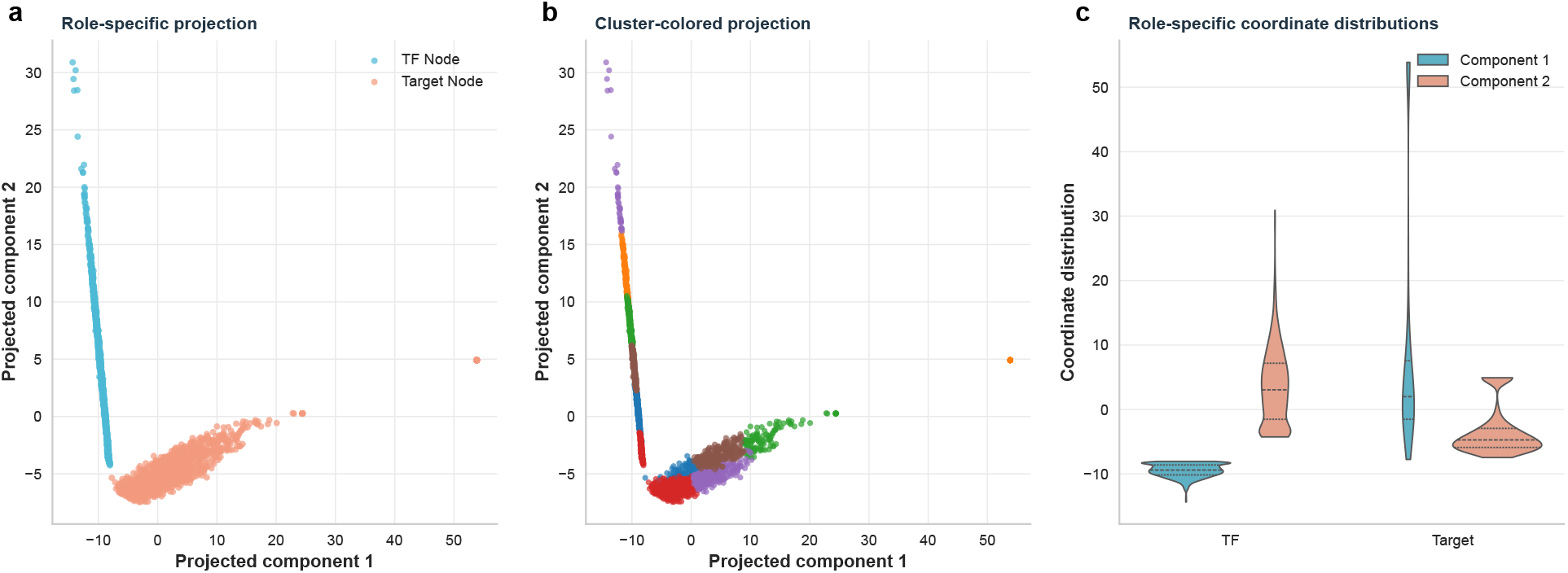
Role-specific latent organization of BRIDGE-GRN embeddings. **a**, Projection of TF-role and target-role embeddings in the representative biological interpretation analysis. **b**, Cluster-colored projection showing the latent organization of the two role-specific embedding spaces. **c**, Coordinate distributions of the projected embedding components, illustrating that the TF-role and target-role representations are related but not interchangeable.

**Figure S12.**
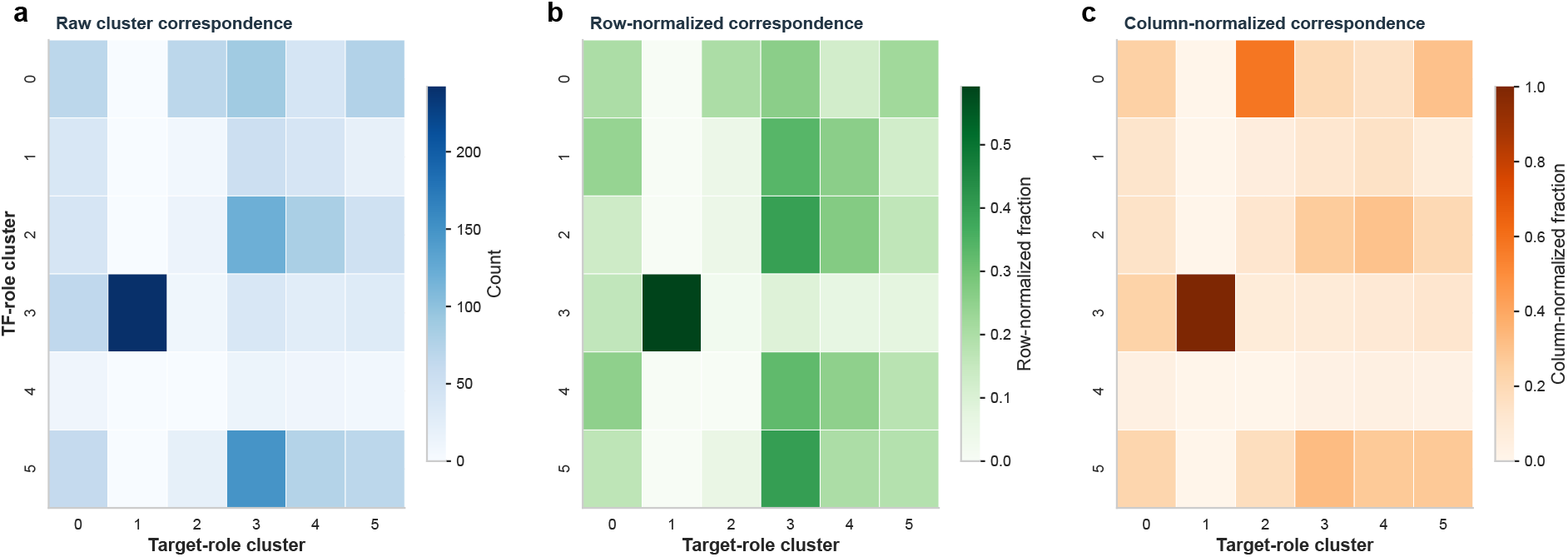
Cluster correspondence between TF-role and target-role embedding spaces. **a**, Raw correspondence counts between TF-role clusters and target-role clusters. **b**, Row-normalized correspondence showing how each TF-role cluster is distributed across target-role clusters. **c**, Column-normalized correspondence showing how each target-role cluster is composed of TF-role clusters.

**Figure S13.**
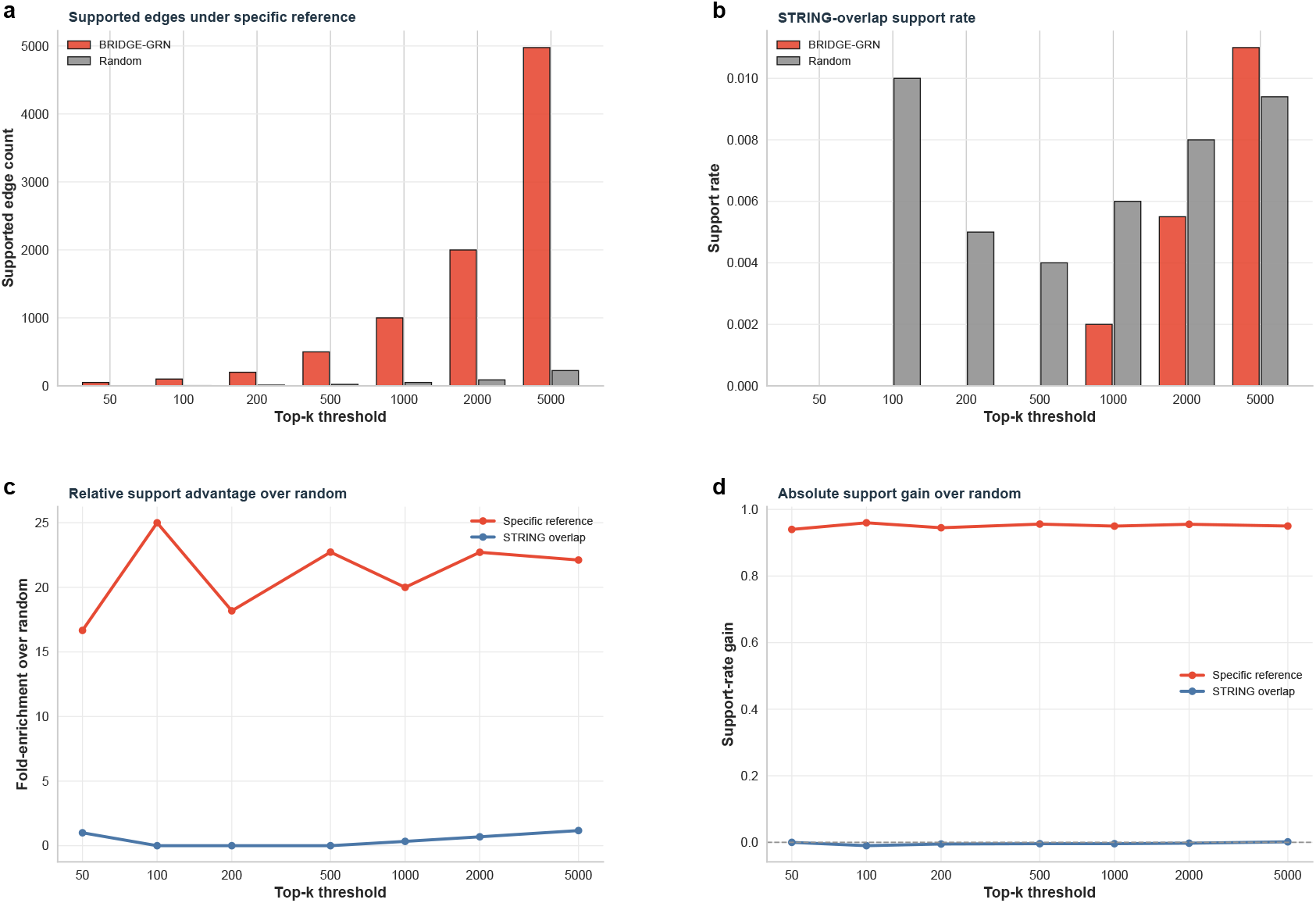
External support enrichment of top-ranked BRIDGE-GRN predictions. **a**, Number of supported edges under the specific-reference support definition across top-*k* thresholds. **b**, STRING-overlap support rate of top-ranked predictions compared with random ranking. **c**, Fold-enrichment of BRIDGE-GRN support over the random baseline under the specific-reference and STRING-overlap definitions. **d**, Absolute support-rate gain of BRIDGE-GRN over the random baseline across top-*k* thresholds.

**Table S14.** Statistical comparisons for ablation experiments. Paired comparisons were conducted across the five matched sample runs. Mean difference is defined as Full BRIDGE-GRN minus the comparison variant. Positive values therefore indicate that the full model achieved higher AUPRC. Confidence intervals are paired 95%confidence intervals.

| Comparison | Metric | Mean difference | 95% CI | *P* value |
| --- | --- | --- | --- | --- |
| Full vs without contrastive regularization | AUPRC | 0.0193 | 0.0121–0.0265 | 0.0017 |
| Full vs single shared tower | AUPRC | 0.0324 | 0.0249–0.0398 | $2.71 \times 10^{-4}$ |
| Full vs identity-view only | AUPRC | 0.0035 | −0.0042–0.0112 | 0.275 |
| Full vs mean fusion instead of concatenation | AUPRC | 0.0069 | −0.0054–0.0192 | 0.195 |
| Full vs smaller embedding dimension | AUPRC | 0.0078 | 0.0032–0.0125 | 0.0094 |
| Full vs bilinear decoder | AUPRC | 0.0038 | −0.0018–0.0094 | 0.134 |
| Full vs cosine decoder | AUPRC | 0.0487 | 0.0436–0.0538 | $1.19 \times 10^{-5}$ |

**Table S15.** Embedding and clustering summary for TF-role and target-role latent spaces. The table summarizes node counts, cluster counts, and projected-coordinate distributions for the role-specific embeddings used in the biological interpretation analysis.

| Role space | Nodes | Clusters | Mean component 1 | SD component 1 | Mean component 2 | SD component 2 |
| --- | --- | --- | --- | --- | --- | --- |
| TF-role | 1620 | 6 | -9.492 | 1.022 | 3.406 | 5.776 |
| Target-role | 1620 | 6 | 9.492 | 19.187 | -3.406 | 3.761 |

**Table S16.** Top-k support enrichment statistics for BRIDGE-GRN predictions and random rankings. Support rates are reported across top-k thresholds under the available external support definitions.

| Method | Support definition | Top-k | Support rate | Fold enrichment |
| --- | --- | --- | --- | --- |
| BRIDGE-GRN | STRING overlap | 50 | 0.0000 | 0.000 |
| BRIDGE-GRN | STRING overlap | 100 | 0.0000 | 0.000 |
| BRIDGE-GRN | STRING overlap | 200 | 0.0000 | 0.000 |
| BRIDGE-GRN | STRING overlap | 500 | 0.0000 | 0.000 |
| BRIDGE-GRN | STRING overlap | 1000 | 0.0020 | 0.240 |
| BRIDGE-GRN | STRING overlap | 2000 | 0.0055 | 0.660 |
| BRIDGE-GRN | STRING overlap | 5000 | 0.0110 | 1.320 |
| Random | STRING overlap | 50 | 0.0000 | 0.000 |
| Random | STRING overlap | 100 | 0.0100 | 1.200 |
| Random | STRING overlap | 200 | 0.0050 | 0.600 |
| Random | STRING overlap | 500 | 0.0040 | 0.480 |
| Random | STRING overlap | 1000 | 0.0060 | 0.720 |
| Random | STRING overlap | 2000 | 0.0080 | 0.960 |
| Random | STRING overlap | 5000 | 0.0094 | 1.128 |
| BRIDGE-GRN | Specific reference | 50 | 1.0000 | 23.773 |
| BRIDGE-GRN | Specific reference | 100 | 1.0000 | 23.773 |
| BRIDGE-GRN | Specific reference | 200 | 1.0000 | 23.773 |
| BRIDGE-GRN | Specific reference | 500 | 1.0000 | 23.773 |
| BRIDGE-GRN | Specific reference | 1000 | 1.0000 | 23.773 |
| BRIDGE-GRN | Specific reference | 2000 | 0.9995 | 23.761 |
| BRIDGE-GRN | Specific reference | 5000 | 0.9952 | 23.659 |
| Random | Specific reference | 50 | 0.0600 | 1.426 |
| Random | Specific reference | 100 | 0.0400 | 0.951 |
| Random | Specific reference | 200 | 0.0550 | 1.308 |
| Random | Specific reference | 500 | 0.0440 | 1.046 |
| Random | Specific reference | 1000 | 0.0500 | 1.189 |
| Random | Specific reference | 2000 | 0.0440 | 1.046 |
| Random | Specific reference | 5000 | 0.0450 | 1.070 |

**Table S17.**
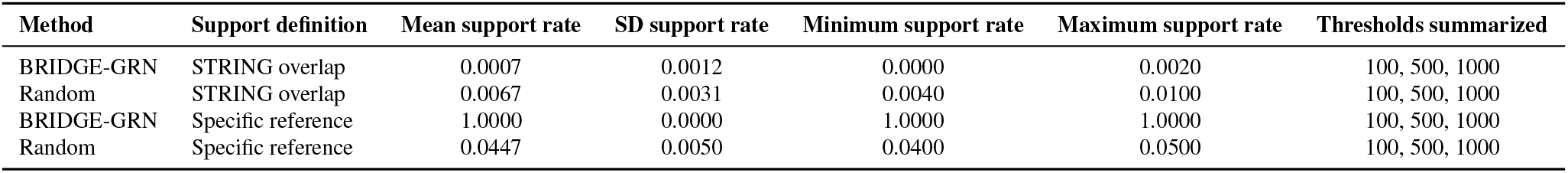
External support summary for top-ranked predictions. Support rates are summarized across three representative top-*k* thresholds, 100, 500, and 1000, for each method and support definition.

**Table S18.** Driver-centered regulatory module summary. The table reports driver ranking scores, supported and candidate outgoing edges, support rates, and mean edge scores for representative driver-centered modules.

| Driver TF | Driver ranking score | Total module edges | Supported edges | Candidate edges | Support rate | Mean edge score |
| --- | --- | --- | --- | --- | --- | --- |
| GABBR1 | 997.329 | 0 | 0 | 0 | — | — |
| NANOG | 724.609 | 8 | 8 | 0 | 1.000 | 0.9996 |
| DR1 | 594.243 | 0 | 0 | 0 | — | — |
| SIN3A | 577.883 | 8 | 8 | 0 | 1.000 | 0.9993 |
| UTF1 | 502.953 | 8 | 8 | 0 | 1.000 | 0.9991 |
| TET1 | 359.506 | 0 | 0 | 0 | — | — |
| ZFP281 | 251.780 | 0 | 0 | 0 | — | — |
| GTF3C1 | 234.986 | 0 | 0 | 0 | — | — |
| MED12 | 174.149 | 0 | 0 | 0 | — | — |
| TCF12 | 159.164 | 0 | 0 | 0 | — | — |
| RNF2 | 133.287 | 0 | 0 | 0 | — | — |
| DDX21 | 106.477 | 0 | 0 | 0 | — | — |
| BTAF1 | 103.463 | 0 | 0 | 0 | — | — |
| NRF1 | 43.730 | 0 | 0 | 0 | — | — |
| POU5F1 | 8.948 | 0 | 0 | 0 | — | — |

**Table S19.**
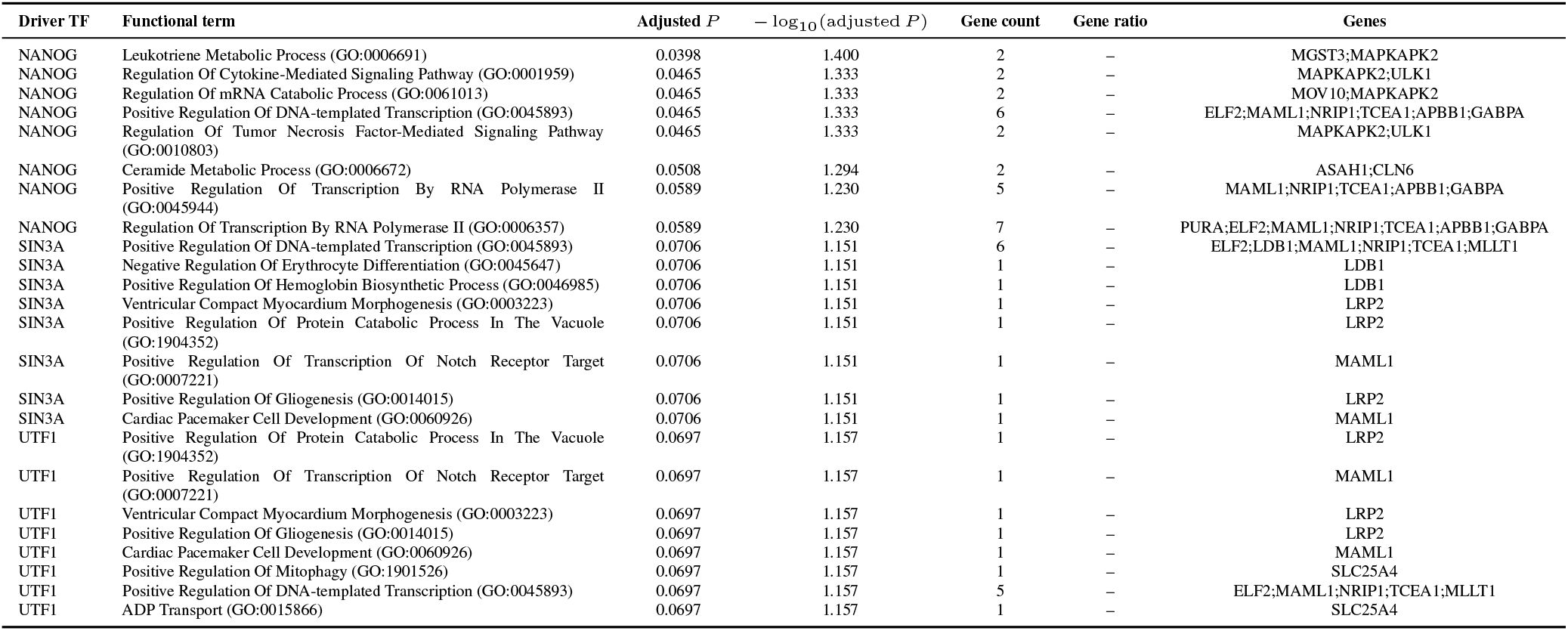
Functional enrichment summary for representative driver-centered regulatory modules. Terms are sorted by adjusted *P* value, with up to eight top terms retained per driver for readability.

